# The Hidden Landscape of Missed Effects in Human Functional Neuroimaging

**DOI:** 10.64898/2026.05.21.726948

**Authors:** Stephanie Noble, Hallee Shearer, Matthew Rosenblatt, Jean Ye, Rongtao Jiang, Link Tejavibulya, Maya L. Foster, Qinghao Liang, Javid Dadashkarimi, Margaret Westwater, Iris Q. Cheng, Max Rolison, Hannah Peterson, Brendan D. Adkinson, Saloni Mehta, Chris Camp, Alexandra Fischbach, Fabricio Cravo, Amanda Mejia, Thomas Nichols, Joshua Curtiss, Dustin Scheinost

**Author notes:** These authors contributed equally to this work (Joshua Curtiss, Dustin Scheinost).

## Abstract

Functional neuroimaging aims to uncover brain processes underlying behavior and disease, yet studies are often underpowered to detect these effects. How this literature has shaped our understanding of brain function remains unknown, and little guidance exists for planning better powered studies. An underappreciated barrier is that commonly reported effect sizes across the brain are inflated, biasing study planning. Here, we introduce a correction for this inflation bias and show how more accurate studies can be planned using corrected effect size benchmarks from a mega-analysis of 63 typical studies across seven large datasets (52,979 participants). We find that common methods of planning studies based on uncorrected effects lead to roughly half the expected detections at typical sample sizes, with limited spatial overlap with original findings. These missed effects collectively explain meaningful additional variance in the desired outcome. We show how to recover missed effects by planning not only for power but also for a target number of detections via corrected benchmarks, or by taking a whole-brain approach with multivariate effects that individual research groups can detect (n < 50 compared to n > 1,000 for a typical univariate effect). These findings lay the groundwork for more informed study planning and a richer understanding of the widespread nature of brain effects, with implications for shared challenges (and solutions) across biomedicine.

## Introduction

Human neuroimaging research has blossomed over the past decades, spurred in part by major international efforts to understand the brain, such as the US BRAIN Initiative and the International Brain Initiative. Functional magnetic resonance imaging (fMRI) efforts in particular have transformed our understanding of brain function in health and disease. However, as the field has matured, mounting evidence has revealed that studies may be underpowered (i.e., have a low probability of detecting effects), resulting in many missed discoveries ^1,2^. Much attention has been paid to one implication of these results: that region-specific findings do not replicate from one study to the next^3,4^. Yet relatively little attention has been paid to how a literature of underpowered studies shapes our current understanding of the nature and spatial extent of brain involvement underlying many psychological processes. Given that fMRI research commonly searches for effects across the brain, understanding the extent of missed effects has important implications for the ongoing debate in neuroimaging (and neuroscience more broadly) about whether the brain is dominated by localized or widely distributed processing^5^.

A central reason for this discrepancy is that while univariate procedures for estimating effect size and power at a single brain area are relatively well established, there are currently no established procedures for quantifying the true mass univariate distribution of effect sizes and power across the brain while accounting for the uncertainty that arises from examining thousands of brain variables simultaneously. A systematic effort is needed to provide more robust effect-size benchmarks to inform power calculations and study planning in functional neuroimaging, and to understand how much of the brain we may be missing. Only recently, with the proliferation of large, publicly shared neuroimaging datasets alongside advances in computational infrastructure for analyzing them, has such a comprehensive investigation become possible.

The present work leverages multiple large-scale neuroimaging datasets, coupled with a new mass-univariate statistical theory, to systematically estimate effect sizes and power across the brain. This work reveals how much of the brain we might be missing when using status quo effect-size estimates that do not account for joint uncertainty, and shows how we can plan more robust studies moving forward. To our knowledge, this represents the first systematic effort to estimate the “true” distribution of effect sizes and power across the brain for common fMRI study designs, thereby establishing empirical foundations for more robust evidence-based study planning and power calculations in the field. This effort not only supports the increased movement towards transparent reporting in human brain mapping, but also connects the field with statistical developments in other biomedical fields facing similar challenges in high-dimensional data (e.g., ^6^).

### Effect sizes across the brain in functional neuroimaging

We performed 63 studies spanning psychological (i.e., cognitive, psychiatric), physical (e.g., age, sex/gender, body mass index; see caveats in *Methods*), and task-based research across seven datasets (total n = 52,979 unique participants) representing diverse populations, study designs, and outcome types (**Fig. 1**; detailed study characteristics in **SI Tables 1-2**). When performing a study, researchers typically estimate “mass univariate” effect sizes for each brain area separately and report the full map of point estimates across the brain (e.g., exemplar studies in **Fig. 1a;** all studies in **SI Figs. 1-2**). However, these mass univariate distributions are inflated—specifically, they exhibit higher variance than the true distribution due to sampling error, narrowing and approaching the true width as sample size increases (cf. **Fig. 1a**). Following in the spirit of a meta-analysis, we developed a principled approach that uses multiple studies to model this decreasing variance and estimate, for the first time, the uninflated distribution of effect sizes across the brain (**Fig. 1c**; *Methods: Estimating population effect size distribution*).

**Figure 1.**
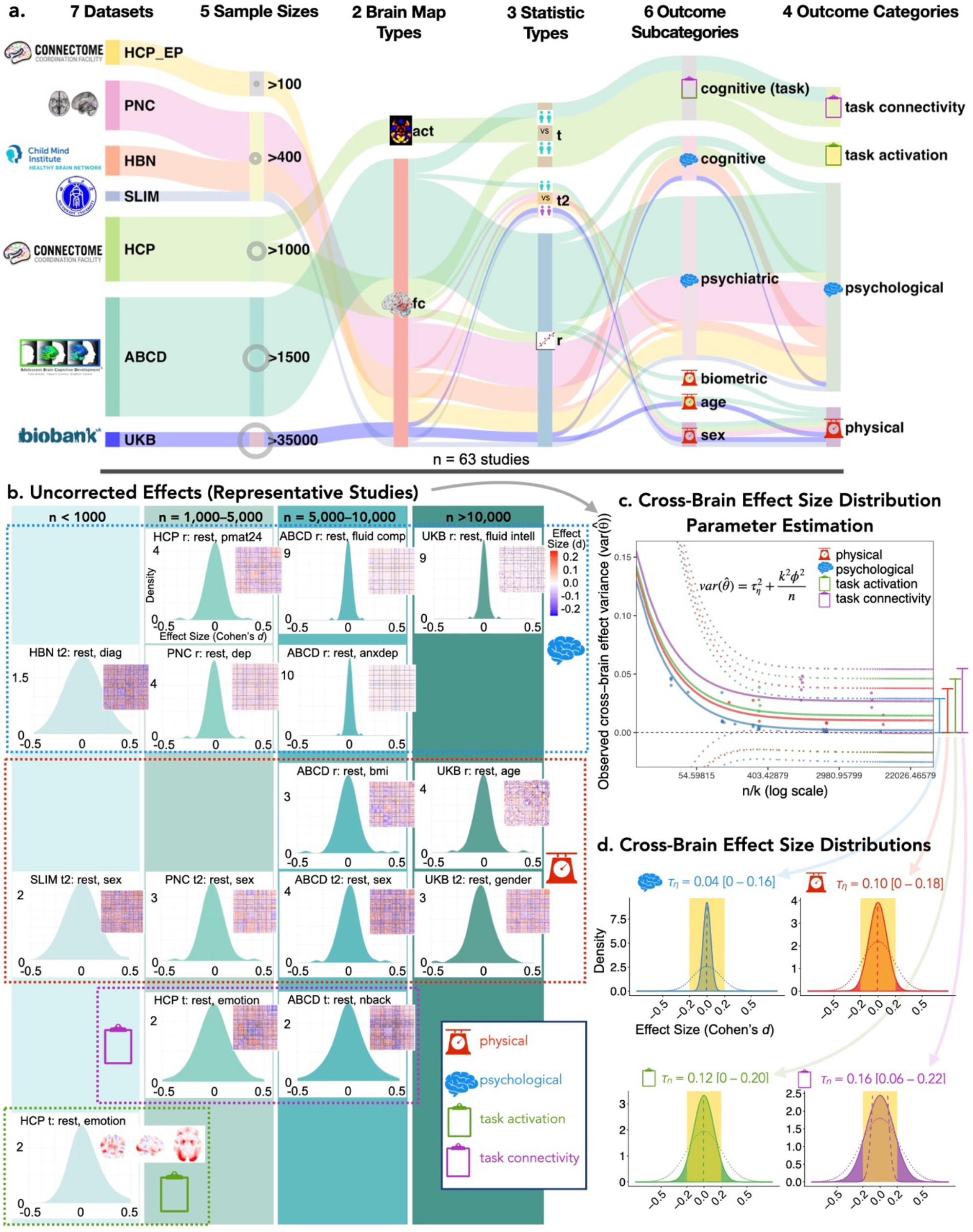
Study characteristics and corrected cross-brain effect size estimation. **(a)** Sankey diagram depicts datasets, sample sizes, brain map types, effect test types, outcome sub-categories, and outcome categories for each study in the database, with the width of each segment of the pathway representing the number of studies in that segment. **(b)** Resulting uncorrected cross-brain effect size distributions for exemplar studies. Density and spatial maps are shown for each study. Studies are arranged by sample size (left to right; also light-to-dark density plot colors) and outcome category (top to bottom boxes). **(c)** Procedure for estimating the distribution of effect sizes across the brain for each outcome category. Parameter estimation plots illustrate how the per-study mass univariate cross-brain variance 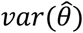 decreases with sample size *n*, asymptotically approaching the true value 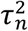 at a rate that depends on the number of test groups *k* (e.g., *k*=1 for 1-sample tests) and within-variable sampling error *ϕ*^2^. Points indicate 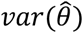 for each study. **(d)** Estimated effect size distributions for each outcome category: psychological (blue), physical (red), task-based activation (green), and task-based connectivity (purple). Density plots are shown for parameter point estimates (solid line) and 95% confidence interval upper and lower bounds (dotted lines). While the point estimate is the best estimate across all study types within a category, the confidence intervals encompass uncertainty about the magnitude of effects across study types. Gold bars indicate effects below the conventional “small” (|*d*|<0.2) effect size threshold. Estimated distributions indicate that most univariate neuroimaging effect sizes fall well below conventional benchmarks. Abbreviations: Adolescent Brain and Cognitive Development: ABCD; Healthy Brain Network: HBN; Human Connectome Project; HCP; Philadelphia Neurodevelopmental Cohort; PNC; UK Biobank: UKB. See **SI Tables 1-2** for detailed study characteristics.

#### Corrected effects are smaller than previously reported

Across all study categories, the majority of effects (79% to >99%) fell below the |*d*| = 0.2 threshold conventionally considered “small” (^7^; **Fig. 1c**; **SI Tables 3-4**). Between-subjects studies, such as brain-behavior associations or group contrasts, exhibited particularly small effects: for a typical study examining physical and psychological variables, 95% and >99% of effects, respectively, fell below |*d*| = 0.2, with only 32% and 1% surpassing the smaller |*d*| = 0.1 mark. Motion correction strategies that did not regress out motion produced modestly larger effects, which may be attributed to residual motion-related confound, even when excluding the highest motion subjects.

These corrected estimates are substantially smaller across study types than previous isolated, uncorrected reports suggest (e.g., ^2,4^), confirming that standard mass-univariate effect-size estimation procedures yield inflated estimates. Researchers designing related studies would benefit from anticipating small effects and planning accordingly.

#### Effect sizes vary across common study types

Psychological associations exhibited the smallest effects, and task-based connectivity exhibited the largest effects, up to four times greater than the former (Fig. 1c). As such, nearly a quarter of task-based connectivity effects exceeded the |d| = 0.2 threshold conventionally denoting small effects, whereas less than 1% of psychological effects exceeded the same threshold. Task-based activation showed the second-largest effects, at 75% the size of task-based connectivity, followed closely by physical effects at 80% the size of task-based activations. Psychological effects showed the biggest difference, being less than half the size of physical effects (40%).

These differences can be largely attributed to two factors. First, the task-based studies represent within-subject designs, whereas the physical and psychological studies used here represent between-subject designs. The larger effect sizes for within-subject studies are partially expected due to the reduced impact of substantial between-subject error. Second, for the within-subject analyses, connectivity analyses use distinct resting-state scans from separate runs to contrast with the task-based scans, whereas activation analyses compare adjacent states within a single scan. Activation maps are also much finer in resolution than connectivity maps, which are based on regions rather than voxels.

Altogether, these findings suggest that researchers should plan for most psychological associations to be very small (explaining the low replicability of cognitive and psychiatric associations previously reported by ^4^), physical associations to be moderately larger but still small, and small-to-medium effects for task-based designs, with the largest effects for connectivity analyses.

#### Larger effects at broader and multivariate scales

Network-level (i.e., pooled within network) and multivariate (i.e., CCA and Hotelling’s T2 test) effects were modestly (network) to substantially (multivariate) stronger than edge- or voxel-level effects (**Fig. 2a**; **SI Tables 3-4**; **SI Fig. 5**). Alongside the need to correct for fewer comparisons, this helps explain the greater power and replicability previously reported for large-scale and multivariate methods (^2,4,8,9^).

**Figure 2.**
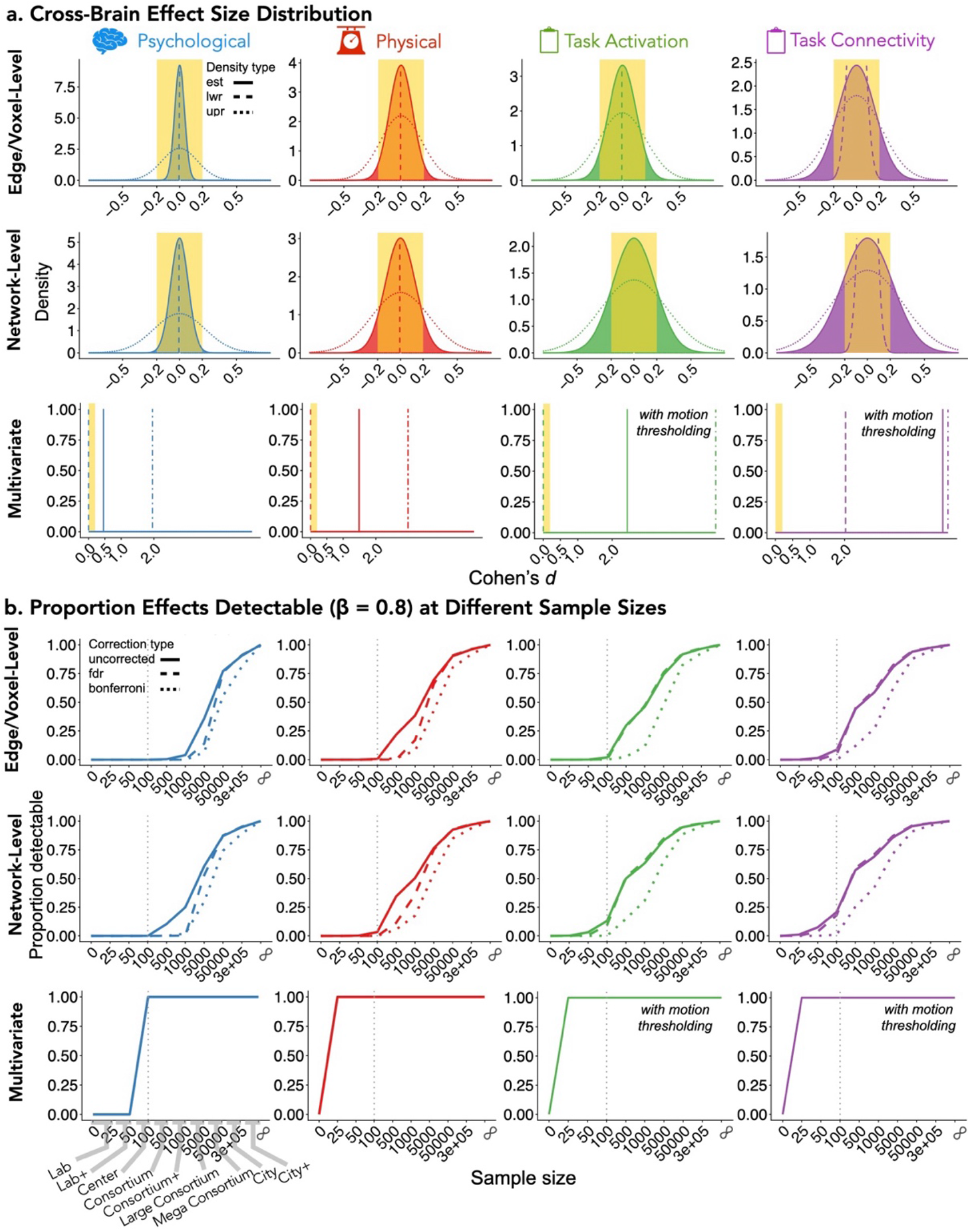
Effect size distributions and statistical power across scales of inference. **a)** Edge- or voxel-level (top), network-level (middle), and multivariate (bottom) estimates of effect size distributions for each outcome category (red, physical; blue, psychological; green, task-based). For multivariate within-subject results only, motion thresholding rather than motion regression was used since the motion regression procedure does not exist for a one-sample multivariate t-test. Gold bars indicate effects below the conventionsal “small” (|*d*|<0.2) effect size threshold. **b)** Sample size required to detect effects with 80% power for edge- or voxel-level (top), network-level (middle), or multivariate (bottom) effects for each outcome category (red and top, physical; blue and middle, psychological; green and bottom, task-based). Each plot shows the proportion of effects across the brain that can be detected with 80% power in each sample size category after Bonferroni, false discovery rate (FDR), or no correction. Sample size categories, chosen to align with observed practices in the field, include: Lab (n = 0–25), Lab+ (n = 26–50), Center (n = 51–100), Consortium (n = 101–500; e.g., HCP-EP; SLIM), Consortium+ (n = 501–1,000; e.g., PNC, HBN), Large Consortium (n = 1,001–5,000; e.g., HCP), Mega Consortium (n = 5,001–50,000; e.g., UKB), City (n = 50,001–300,000), and City+ (n = 300,000+). A grey dotted line denotes sample sizes expected to be attainable at individual sites versus those requiring multiple sites (i.e., Consortium).

### Study planning implications using corrected effect size benchmarks

Having established corrected effect size benchmarks, we next performed a mass univariate power analysis to illustrate sample size requirements for robust study planning (e.g., ^10,11^; **Fig. 2b**). To accomplish this, we developed a framework to estimate mass univariate power given a cross-brain distribution of effect sizes under different multiple testing control procedures including Bonferroni, false discovery rate (FDR), and no control settings (see *SI Methods: Estimating cross-distribution power with multiple testing correction*).

At the edge- and voxel-levels, the vast majority of effects could not be detected with adequate power (β = 80%) at less than Consortium-scale sample sizes (n ≤ 100 subjects). Between-subjects studies required Large or Mega Consortium-scale sample sizes to detect more than half of all brain effects, even without multiple-comparisons correction (n ≅ 16,000 for psychological effects; n ≅ 2,000 for physical effects). Task-based effects were more detectable; detecting half of all effects required approximately half the sample size for task-based activation compared with physical outcomes (e.g., n ≈ 1,500 vs. n ≅ 2,000 uncorrected; n ≅ 1,500 vs. n ≅ 3,500 with FDR; n ≅ 5,000 vs. n ≅ 13,000 with Bonferroni) and fewer still for task-based connectivity (e.g., n ≅ 700 uncorrected; n ≅ 700 with FDR; n ≅ 3,500 with Bonferroni). Some task-based connectivity effects were even detectable at sample sizes attainable within individual sites (n ≤ 100), reflecting the power advantages of within-subject designs (**SI Table 5**).

Power increased substantially with broader-scale approaches. At the network level, a non-negligible proportion of task-based effects were detectable at the Center scale with FDR correction (∼5%–15%). More than a quarter of physical effects became detectable at the Consortium+ scale with FDR correction, though psychological effects still required Large Consortium sample sizes. Critically, all multivariate effects were detectable at sample sizes within reach of individual labs^12^ for physical and task-based studies (n ≤ 25) or individual sites for psychological studies (n ≤ 100).

Power varied as expected across multiple testing approaches (i.e., uncorrected > FDR > Bonferroni), although FDR correction approached power levels similar to no correction for n > 100 task studies, underscoring its practical utility. Motion correction strategies that did not use regression resulted in a moderate increase in power, again attributed to relatively strong motion-related effects (**SI Fig. 6-7**).

Overall, the corrected benchmarks indicate that detecting univariate between-subject effects requires consortium-scale samples, aligning with and extending prior estimates for replicating results^4^. However, the situation improves considerably for task-based studies. Broader-scale and multivariate approaches substantially reduce requirements, making adequately powered studies possible with sample sizes within reach of individual labs or sites.

### Consequences of study planning based on uncorrected effects

Whereas the above power analyses are based on new corrected benchmarks, researchers currently rely on uncorrected effects when planning studies. To evaluate how this biases results, we simulated studies planned from uncorrected “basis” studies and compared them against studies planned from corrected benchmarks. Uncorrected strategies mirror common practices when planning based on a published or pilot study. The “optimistic literature” and “conservative literature” strategies used maximum and minimum significant effects respectively, reflecting the fact that publications typically report only significant effects (with known selection bias^1^). The “ideal pilot” strategy used the mean unthresholded effect, reflecting a scenario where the researcher has access to the full effect map (e.g., a pilot study) and conservatively targets a typical rather than stronger effect. Basis and main studies were simulated across 4 outcome categories, 6 sample sizes, and 3 planning strategies (200 repetitions each; 28,800 total simulated studies), with an additional 4,800 simulated studies planned directly from corrected benchmarks.

Using an uncorrected basis study yielded unpredictable results that generally failed to capture expected findings. When basis studies used sample sizes within reach of individual labs (n ≤ 100), main studies often detected far fewer true positives than anticipated (**Fig. 3a**). The “pilot” strategy that avoids selection bias improved expectations, yet even with basis studies of 50 subjects and main studies featuring hundreds of subjects, all but the task connectivity condition still missed at least a quarter of expected effects. Furthermore, the “pilot” strategy missed many original effects at most sample sizes, with recovery peaking at a point where the main study sample size dwarfs the basis study and then steadily decreasing as the gap narrows and reverses (**Fig. 3a**, grey plots). At planned sample sizes representing hundreds of thousands of dollars in investment, the practical cost of so many missed expectations is high.

**Figure 3.**
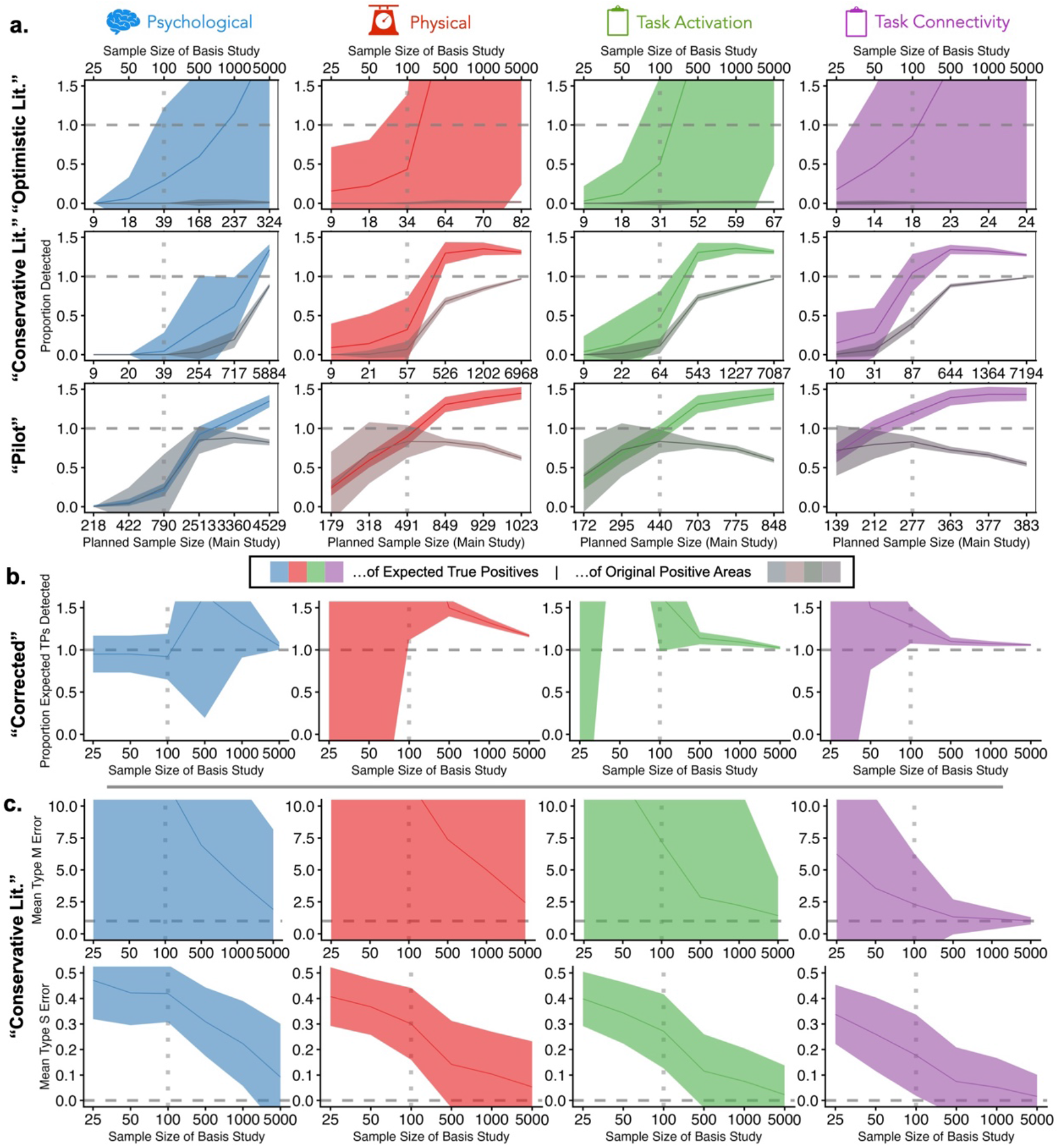
Fewer detections than expected when planning studies using uncorrected effects. **(a)** Simulations show expected compared with actual results when planning studies to replicate the results of another uncorrected study (e.g., pilot or published study), whether using the maximum significant effect (top, “Optimistic Literature” strategy), minimum significant effect (middle, “Conservative Literature” strategy), or mean positive effect (bottom, “Pilot” strategy) obtained from the basis study. Saturated ribbon plots show proportion of expected effects actually detected (true positives). Unsaturated (more gray) ribbon plots show proportion of original findings recovered. **(b)** Proportion of expected true positives actually detected when planning studies based on corrected effects. **(c)** Type M (magnitude) and S (sign) errors for the “Conservative Literature” strategy. Lines and ribbon widths reflect mean and standard deviations over 200 repetitions. Dashed horizontal lines denote target levels (above is better for **(a)** and **(b)** and below is better for **(c)**), and dotted vertical lines denote sample sizes within reach of individual labs (below is better).

In contrast, planning from corrected effect size estimates yielded true positive detection rates that met or exceeded expectations (**Fig. 3b**). Expectations seem fairly conservative at n < 500, but this represents a small difference where only a couple effects are expected and dozens are detected, so researchers can be confident in not missing effects. Corrected benchmarks and theory also offer a new study planning approach: the ability to explicitly target an expected number of detections alongside a level of statistical power, rather than treating the spatial extent of missed effects as an afterthought.

### Missing effects add meaningful explanatory power

Having established that typical studies detect only a fraction of all contributing brain areas, we next examined whether those undetected areas contribute meaningfully to the outcome of interest. In light of the idea that the null hypothesis is typically implausible in psychology (i.e., given enough data, a test will be significant, even if effects are exceedingly small), this moves the target to understanding whether the addition of brain areas outside of a more focal set of regions adds more than a trivial contribution to the story. To address this, we estimated the multivariate effect size as progressively more brain areas were included in the analysis, selecting areas in order of univariate effect size to mirror standard practice of reporting the most significant regions (**Fig. 4**). After univariate feature selection, we again used PCA combined with CCA to obtain stable multivariate estimates (see *Methods*) that reflect the maximal combined effect size contributed across all included variables.

**Figure 4.**
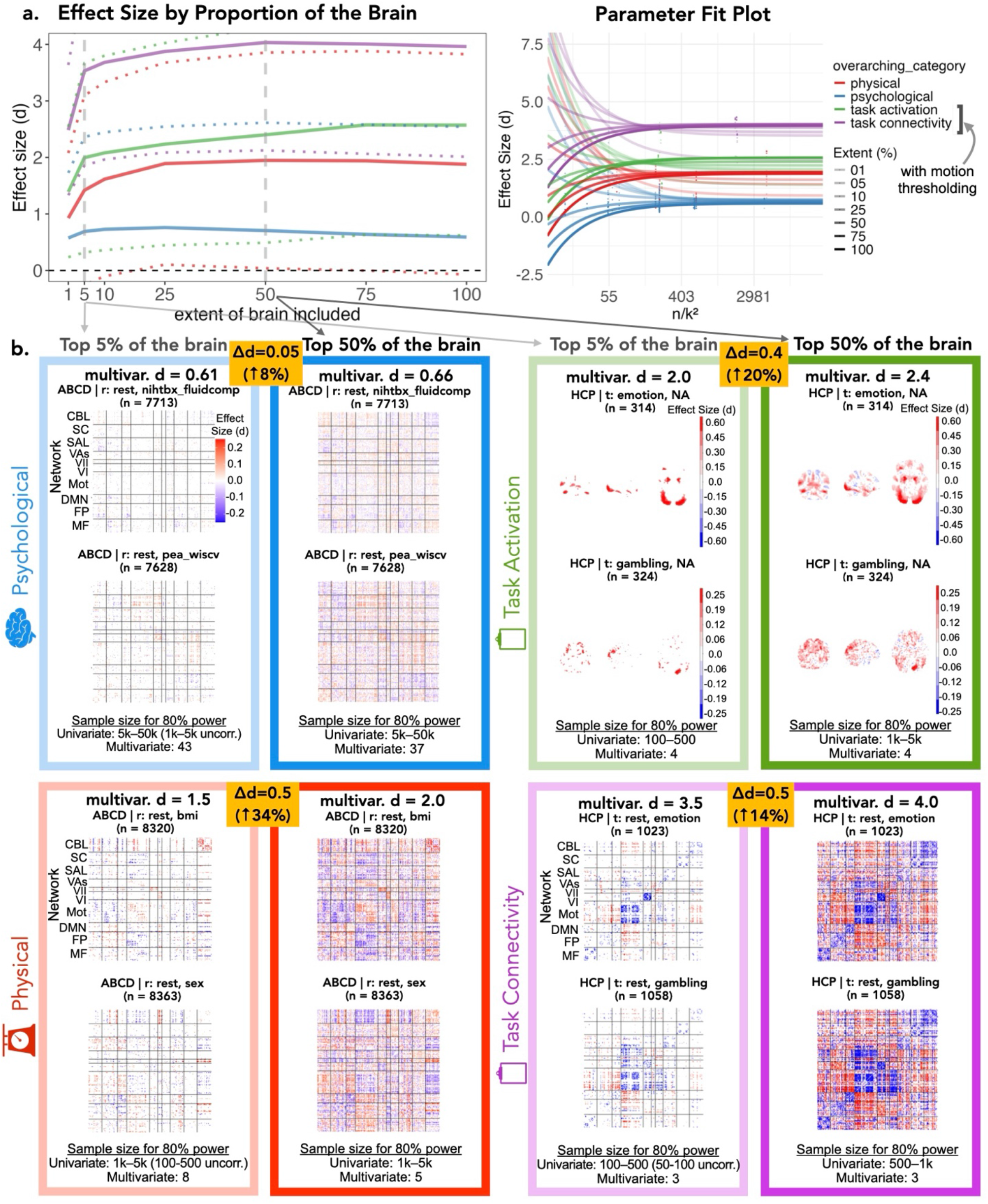
Effect size increases with the proportion of the brain included. **Top panel: a)** Multivariate effect size from the top *k* strongest effects, by outcome category (red, physical; blue, psychological; green, task activation; purple, task connectivity). The top 1%, 5%, 10%, 25%, 50%, 75%, and 100% marginal effects (x-axis) were selected and used to estimate the multivariate effect size. The parameter estimation plot for each top *k* effect is shown to the right, with higher k in more opaque colors. Dotted lines denote top 5% and top 50% of effects. **b)**. Multivariate and univariate results for each category at top 5% of areas (left, lighter box) and top 50% of areas (right, darker box). Spatial plots show univariate effect sizes (uncorrected) from representative studies. Multivariate effect sizes across all included areas are provided at the top of each box. Bottom text shows sample sizes needed to obtain adequate (80%) power on average for the top *k* univariate effects, and sample sizes needed for adequate power for the multivariate effect size (estimated parametrically). Spatial maps are arranged by canonical networks (functional connectivity maps) or spatial location (activation maps). Canonical brain networks include MF: medial frontal, FP: frontoparietal, DMN: default mode network, Mot: motor, Vas: Visual Association, SAL: salience, SC: subcortex, CBL: cerebellum. Brighter red indicates larger positive effects; brighter blue indicates larger negative effects.

While the top 5% of brain areas contributed more than half of the maximum effect size, effects increased meaningfully when including the top 50% of brain areas. Physical effects increased the most, increasing from *d*=1.5 with 5% of the brain to *d*=2.0 with 50% of the brain (34% increase). Task-based effects increased by a moderate amount, with activation increasing by 20% and connectivity by 14%. Psychological outcomes showed the smallest increase from *d*=0.61 to *d*=0.66 (8% increase) when increasing from 5% to 50% of the brain. This is complicated by the fact that the brain accounts for such a small overall variance, meaning that the addition of signal with more brain areas may compete with the addition of substantial noise.

Altogether, this work points to non-trivial additional variance explained by including at least 50% of the brain, unless the maximum explainable variance is already low. One notable exception arose without motion regression: at a certain point, adding brain regions sometimes reduced rather than increased variance explained, particularly for psychological (peak at ∼10% of the brain) and physical (peak at ∼50%) outcomes under motion thresholding. This likely reflects the dominating influence of noise affecting components obtained during dimensionality reduction when high-motion artifact is not fully removed (**SI Fig. 8**).

### Turning anecdote into evidence for study planning

In the functional neuroimaging community, it is a common refrain that studies are often underpowered (cf. ^3,12,13^), yet effect size and power benchmarks have proven difficult to establish, obscuring the spatial extent of this problem. The corrected theory and benchmarks introduced here reveal a literature that systematically underestimates the landscape of detectable effects, and that missed effects are not trivial. Overall, these findings invite researchers to reconsider the implications of a literature that overlooks many meaningful effects and to embrace study planning methods better suited to capturing subtle effects across the brain.

#### Published findings are only the tip of the iceberg

In the absence of empirically grounded guidelines, the neuroimaging literature has been historically shaped by underpowered studies that we show are expected to miss many non-trivial brain effects. This collective knowledge only represents the tip of the iceberg.

Some evidence has pointed towards more spatially extensive brain involvement in psychological processes than is widely recognized (cf. ^5^), but the degree to which missed areas are meaningful enough to warrant moving beyond focal approaches has remained unaddressed. As expected, a small portion of regions contribute more than others; the top 5% of brain areas collectively capture more than two-thirds of the maximum variance explainable by the whole brain. However, the non-trivial improvement from including more of the brain (∼25% increase unless exceedingly small) reveals that a focal perspective captures only part of the story.

These findings shift the focus from whether effects are present—which contemporary perspectives in statistics consider a largely trivial question underlying null hypothesis significance testing (e.g., ^13^)—towards how many effects are meaningful or how much of the brain should be considered to characterize a phenomenon. Any threshold will, by construction, exclude regions that do contribute statistically meaningful additional explained variance. Whether that line should be drawn based on meaningful added contribution (e.g., is a 10% increase in variance explained by increasing from 50% of 70% of the brain worth it for task activation?) or on total variance explained (e.g., is 75% variance explained from 5% of the brain sufficient for characterizing task-based differences in connectivity?) is an important question for future work, which will require careful characterization and development of consensus to motivate what constitutes a meaningful effect.

#### Planning conservatively for future studies

These findings invite researchers to rethink reliance on conventional study planning procedures and provide a concrete touchstone to ground expectations moving forward. While the corrected benchmarks presented here largely support the expectation of small univariate effect sizes for most study types, the situation is not monolithic. Within-subject task-based designs offer substantially larger effects than between-subjects designs, to the extent that researchers may be able to conduct adequately powered studies at sample sizes attainable within individual labs. For the many researchers conducting between-subjects studies, however, the corrected benchmarks underscore the need for either large collaborative datasets or more powerful inferential approaches.

Multivariate and network-level approaches represent a practical route to adequately powered studies across study types, offering detectable effects at sample sizes within reach of individual labs while also being more reliable and replicable^4,8,9,14^. This benefit is due in part to the reduced burden of multiple testing relative to edge- or voxel-level approaches. Thus, broader-scale approaches offer not only theoretical but also pragmatic advantages.

The statistical procedures developed here derive from basic principles of sampling error that we extended to characterize effect-size distributions and power in high-dimensional neuroimaging data. This strategy avoids the limitations of relying either on marginal estimates from simultaneous confidence intervals (too uncertain given the high dimensionality) or on point estimates from individual studies (inflated due to sampling variability; cf. *Supplementary Methods*). To our knowledge, these corrected distributions provide the first empirical whole-brain benchmarks for study planning in neuroimaging. For these or any other effect size estimates, we recommend that researchers consider both the main parameter estimates and their uncertainty when planning studies^15,16^.

Although revision is warranted, the situation is not as dire as it may seem. Brain imaging is a relatively new and remarkable field, and, in the prescient words of Jacob Cohen, “In new areas of research inquiry, effect sizes are likely to be small (when they are not zero!) [because they are] typically not under good experimental or measurement control or both”^7^. Indeed, best practices for measurement, acquisition, and analysis are continuously being refined for neuroimaging in the face of its complex and historically unwieldy data characteristics, and patience is likewise warranted when studying such subtle, individualized phenomena as mental life^5^. The scale of neuroimaging discoveries that are replicable remains impressive (e.g., the ability to map critical language cortex during surgery, the discovery of classes of large-scale brain networks), especially keeping in mind that even relatively well-characterized phenomena exhibit “small” effect sizes (e.g., the difference in IQ similarity between twins and non-twins; the height difference between 15 and 16 year old girls^7^).

#### Limitations

This work represents a first step toward a comprehensive database of standard effect sizes for fMRI research, beginning with typical study designs from some of the largest currently available datasets. Several extensions to this resource would substantially broaden its scope and utility.

First, generalizability is currently limited to the study categories included here. These benchmarks would benefit from inclusion of additional task-based designs, within-subject contrasts, longitudinal designs, scan protocol variations (including scan length; cf. ^18^), and various processing decisions known to substantially impact results^3,17^, particularly timeseries-level artifact attenuation strategies. Population-specific differences between datasets also warrant future investigation.

There is also a wealth of untapped data that can be integrated into smaller-scale studies. Repositories such as NeuroVault^19^ could be used to include studies that represent a broader range of research contexts than the select large datasets included here. However, aggregating numerous smaller studies presents unique scalability challenges. The contribution format introduced previously^20^ offers a starting point, but alignment with existing standards^21^ and automated study labeling approaches will be needed. These challenges are shared with publication-based meta-analysis tools BrainMap^22^ and NeuroSynth^23,^ as well as other study aggregation projects, and represent a timely opportunity to leverage advances in generative artificial intelligence.

Finally, we modeled the effect size distribution parameters by estimating how they would change with sample size as the sample size increased, but this approach may not generalize across all neuroimaging contexts. Additional work is needed to characterize and improve precision across diverse study types and populations. The present work quantifies univariate, network-level, and multivariate power, but does not comprehensively address power for fMRI-specific inferential approaches such as cluster-based inference (cf. ^24^ for power estimation in HCP task activation data), although we note that FDR correction can be more powerful than cluster-based inference^2^. To support future methodological developments, we have made the full set of effect sizes and estimation procedures publicly available.

#### Future Directions

Effect size benchmarks are not only essential for informing future neuroimaging research but also for contextualizing findings and sharing solutions across medical disciplines. For example, genetic studies have shown similarly small effects across millions of common genetic loci, requiring hundreds of thousands to millions of subjects to detect^25^. The field of statistical genetics has spent more than a decade developing methods to estimate the distribution of effects in high-dimensional data while accounting for widespread, correlated effects^6^. While the extent to which genomic assumptions apply to neuroimaging remains unclear (e.g., linkage disequilibrium, sparse causal factors), statistical genetics has navigated many of the methodological challenges neuroimaging is now facing and offers a valuable guide (cf. ^25,26^). Behavioral psychology also offers some lessons in revising expectations and appreciating the implications of likely small effects (cf. ^16,27^; although effect sizes have been shown to be an order of magnitude larger, reports are also considered inflated and thus effects may be quite small^27,28^).

Beyond informing traditional study planning, this work is intended to support methodological advances in neuroimaging. We previously introduced BrainEffeX^20^, an interactive app for exploring the study-level effect size estimates underlying this work (e.g., **SI Fig. 2, 4**; again noting that the uncorrected estimates should not be used directly for study planning). Building on the corrected benchmarks established here, PRISME^29^ and the forthcoming BrainPowerX web app (R00MH130894) together provide resources for empirically informed study planning. The long-term success of these and a wide variety of related open science efforts championed by the neuroimaging community critically depends on sustained public investment in the development, maintenance, and dissemination of methods.

This work aligns with a growing trend in neuroimaging towards leveraging ever-larger datasets and computational resources to revisit the foundational assumptions that underlie modern research practices. While the conceptual core may seem simple, realizing it required new statistical theory and a mega-analysis spanning 52,979 participants—a scale that has only recently become possible through widespread collaborative and data-sharing efforts. For a field grappling with well-documented reproducibility challenges, the shift from convention-based to empirically grounded study planning marks a critical transition: it alters the questions researchers can ask, what they might expect to find, and, ultimately, what we can learn about how the brain functions in health and disease.

## Methods

### Datasets

Seven large-scale datasets (n = 100–40,000; 52,979 total participants) were selected based on the availability of functional neuroimaging data and sample sizes >100 participants. Datasets spanned developmental stages from childhood and adolescence (PNC, ABCD, HBN) to young adulthood (SLIM, HCP, HCP-EP) and mid-to-older adulthood (UKB), including both typically developing populations and clinical cohorts. Imaging protocols included resting-state and task-based fMRI with diverse phenotypic assessments (**Figure 1**; detailed study characteristics in **Supplementary Table 1** and *Supplementary Methods*).

### Preprocessing and subject-level test statistics

*ABCD, HBN*, and *PNC*: Structural images were skull-stripped (FSL), and functional images were motion- and slicetime-corrected (SPM8). Subsequent steps occurred in BioImage Suite, including nonlinear registration of structural images to MNI space and linear registration of functional images to structural images. Noise covariates were regressed, including linear and quadratic drift, a 24-parameter motion model, mean cerebrospinal fluid (CSF), white matter (WM), and global signal (GS). Data were temporally filtered (Gaussian cutoff ≈ 0.12Hz), and functional connectivity was estimated as Fisher z-transformed Pearson correlations between mean timecourses for Shen region pairs.

#### HCP connectivity

Minimally preprocessed data were obtained from the HCP repository, which included gradient correction, motion correction, fieldmap correction, and linear registration. Subsequent processing in BioImage Suite included nuisance regression (linear/quadratic drift, 24-parameter motion model, CSF/WM/GS), temporal filtering (Gaussian cutoff ≈ 0.12Hz), and nonlinear registration to MNI. Functional connectivity was calculated as above, with right-left and left-right phase-encoding connectomes computed separately and then averaged.

#### HCP activation

Beta coefficient maps were obtained from the repository. Volume-based preprocessing included minimal preprocessing, MNI nonlinear registration (FNIRT), intensity normalization, and smoothing (4mm FWHM). Activations were estimated via general linear modeling with task predictors (canonical HRF), temporal derivatives as confounds, high-pass filtering (200s cutoff), and prewhitening.

#### SLIM

Connectivity matrices were obtained from the repository. Processing included discarding the first 10 volumes, slice-timing correction, motion correction, normalization, resampling to 3mm, smoothing (6mm FWHM), linear detrending, bandpass filtering (0.01-0.08Hz), and nuisance regression (six motion parameters, mean CSF/WM/GS) using DPARSF.

#### UKB

Connectivity matrices were obtained from the repository. Processing included MNI152 warping (FNIRT), tissue segmentation (FAST), motion correction (MCFLIRT), intensity normalization, highpass filtering (σ=50s), and artifact removal (ICA-FIX). Group-ICA (dimensionality 25) produced network nodes for dual-regression based connectivity, calculating correlations between node timecourses.

### Group-level test statistics

Subject-level maps were analyzed using three statistical tests: a one-sample t-test, a two-sample t-test, and a correlation test. Three motion adjustment strategies were evaluated: no correction, statistical control (including motion as a predictor), and thresholding (excluding subjects with > 0.1 mm mean frame-to-frame displacement, previously shown to exclude the top 5% of high-motion subjects in the HCP; unpublished). Three scales of inference were examined: edge-/voxel-level (mass univariate), network-level (pooled within Shen 10-network atlas), and multivariate whole-brain level, which used Principal Component Analysis (PCA) to reduce data to the top *k* elements (i.e., k = sample size / 50) followed by Canonical Correlation Analysis (CCA) to obtain the top canonical correlation or Hotelling’s T^2^ test statistic. The PCA step is used to obtain stable estimates^30^. Analyses resulted in mass univariate or multivariate t-statistics or correlation coefficients.

### Effect size conversion

Test statistics were converted to Cohen’s *d* following standard conventions^7^. For the one-sample t-test, 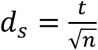 and 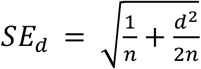 . For the two-sample t-test, 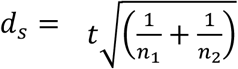 and 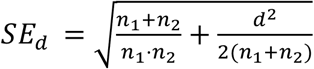 . For correlation, 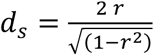 (cf. ^7^) and 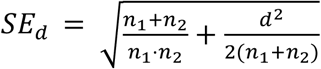, taking group sizes to be equal (i.e, *n*_1_ = *n*_2_ = *n*/2). Multivariate effects were also converted to *d* following the above conversions.

### Estimating cross-brain effect size distributions

To correct for magnitude inflation in the cross-brain effect size distribution due to sampling error, we modeled how observed variability in effect sizes changes with sample size. We formulated a relationship between the observed (uncorrected, mass univariate) cross-brain variance 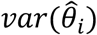 and true cross-brain variance *τ* _*η*_^2^as

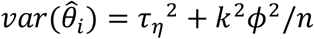

where *ϕ*^2^ is the population variance, *k* is the number of groups for the test, and *n* is the total sample size for the study. Within each outcome category, we estimated *τ* _*η*_ ^2^ and *ϕ*^2^ as the coefficients of

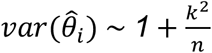

using meta-regression with random intercepts for the dataset/outcome category. For zero-centered normal distributions, the expected magnitude is given by the CDF for the half normal 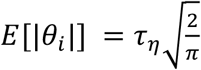, with 68% of effects falling below *τ* _*η*_ in magnitude.

### Power for a distribution of effect sizes with multiple testing correction

Given a cross-brain distribution of effect sizes, the expected number of effects surpassing a target power threshold and average power were derived as

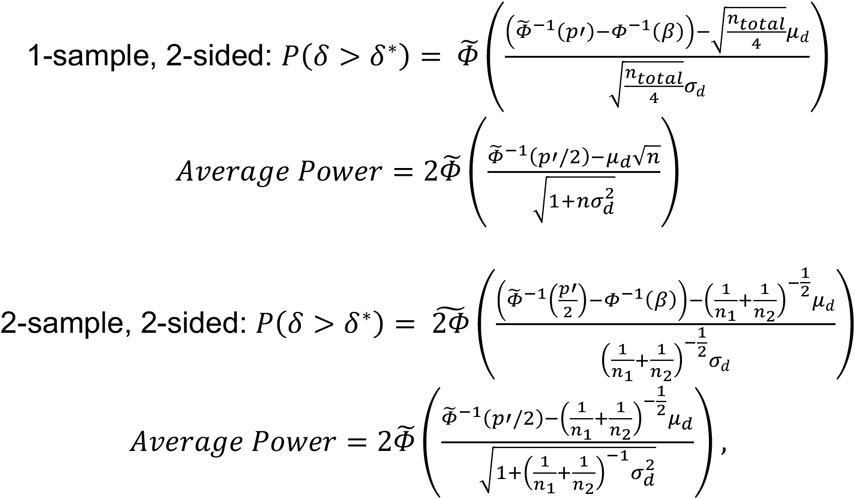

where *n*_*total*_ is the total sample size, *n*_1_ and *n*_2_ are the sample sizes for each group, *p′* is the p-value threshold for rejecting the null, *β* is the target power level, 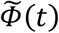 is the standard normal complementary CDF, and effect sizes are normally distributed with mean *μ*_*d*_ and variance 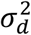 . For an uncorrected test, we use *p*′ = 0.05 and, for Bonferroni correction, *p*′ = 0.05/*m*, given *m* variables. For false discovery rate (FDR) control, *p*′ depends on the data; thus, we derive the following expression for the FDR power implicit function at level *α*_*FDR*_ (here, *α*_*FDR*_ = 0.05) for a 2-sided test as 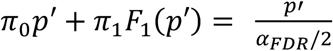, and the alternate p-value distribution (*F* ) for 1- and 2-sample tests as

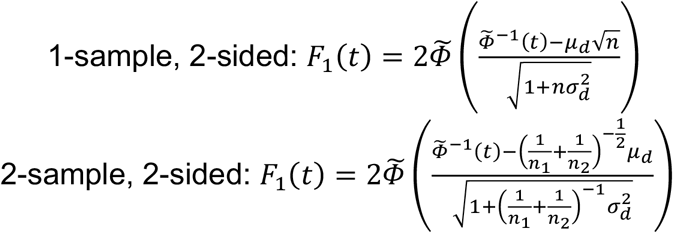

### Full derivations available in *Supplementary Methods*

#### Simulated studies based on corrected and uncorrected effect sizes

For a given outcome category, ground truth effect sizes for 500 brain regions were drawn from *N*(0, *τ*_*η*_^2^) (level 2) and subject-specific error for 10,000 subjects was drawn from *N*(0, *ϕ*^2^) (level 1), yielding a master population with observed effect sizes for each region.

The following steps were conducted for four study planning strategies (“basis sample size,” “basis max effect,” “basis average effect,” and “corrected effect”) at each of six *a priori* sample sizes *n* = [25, 50, 100, 500, 1,000, 5,000].

For strategies using basis studies, a basis study was conducted by drawing the *a priori* sample size from the population and performing a one-sided, one-sample t-test. The main study sample size and researcher-expected true positives (*TP*_*E*_) were defined per strategy as follows. For the *“optimistic literature”* strategy, sample size was calculated to obtain adequate power for the maximum significant effect via conventional power analysis (i.e., pwr.t.test), and *TP*_*E*_ = 1 × *β*, where *β* is target power. For *“conservative literature,”* sample size was calculated to obtain adequate power for the minimum significant effect via conventional power analysis (i.e., pwr.t.test), and *TP*_*E*_ = 1 × *β*. For *“pilot”*, sample size was calculated to obtain adequate power for the mean of the 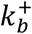 positive-signed effects in the basis study (no significance-based selection), and 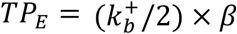 . The main study was then conducted, true positives defined as detections matching the ground truth direction (*x*_1|1_, following notation in ^2^), and researcher expectation errors evaluated as follows:

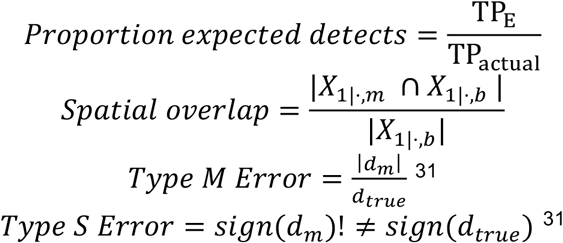

For the strategy using corrected effects, *TP*_*E*_ = *Average Power*_*FDR*_ × *k*, where average power is defined above and *k* is the number of regions. The main study was run at the *a priori* sample size, with true positives and expectation errors defined as above. Only spatial overlap is undefined since no *a priori* spatial location is specified.

This produced 33,600 total simulated studies (200 repetitions × 4 outcome categories × 6 sample sizes × 7 study types reflecting 3 basis, 4 main).

#### Effect size by proportion of the brain

The same procedure used previously to estimate multivariate effect sizes was again used for this analysis (i.e., PCA followed by CCA or Hotelling’s T-squared test and conversion to *d*), with the exception that first the largest *k* percent absolute value effect size predictors were retained before multivariate effect sizes were estimated, across values of *k* from 1% to 100% of the brain. This reflects the size of the effect due to all *k* effects in combination.

## Data availability

All raw data used in the present study have been made publicly available by the contributing repositories in accordance with data use and access regulations set by the respective consortia, as summarized below. The Northeastern University and Yale University Human Research Protection Programs approved secondary analyses of these datasets.

Human Connectome Project (HCP) and Human Connectome Project Early Psychosis (HCP-EP) data are available through the HCP repository (https://www.humanconnectome.org/study/hcp-young-adult ; https://www.humanconnectome.org/study/hcp-early-psychosis). Users must agree to data use terms before accessing ConnectomeDB; details are provided at https://www.humanconnectome.org/study/hcp-young-adult/data-use-terms. UK Biobank data are available to approved researchers conducting health-related research in the public interest through https://www.ukbiobank.ac.uk/. This research was conducted using the UK Biobank Resource under application number 140089. Adolescent Brain and Cognitive Development (ABCD) data are available through the NIH Brain Development Cohorts (NBDC) Data Hub (https://data.NBDC.nih.gov/). The ABCD data used in this report came from Release 2.0.1, NDA Study DOI: 10.15154/1504041. Philadelphia Neurodevelopmental Cohort (PNC) data are available through dbGaP under accession phs000607.v1.p1 (https://www.ncbi.nlm.nih.gov/projects/gap/cgi-bin/study.cgi?study_id=phs000607.v1.p1) for authorized researchers with approved Data Use Certification. Healthy Brain Network (HBN) data are available through the Child Mind Institute LORIS database (http://fcon_1000.projects.nitrc.org/indi/cmi_healthy_brain_network/) for approved researchers. Southwest University Longitudinal Imaging Multimodal (SLIM) data are available under Creative Commons Attribution-NonCommercial license through the International Data-sharing Initiative (https://fcon_1000.projects.nitrc.org/indi/retro/southwestuni_qiu_index.html).

## Code availability

All code used for running statistical analyses has been made publicly available on GitHub (https://github.com/neuroprismlab/calculate_effeX ; https://github.com/neuroprismlab/BrainEffeX_utils ; https://github.com/neuroprismlab/crossbrain_effects). Initial group-level statistics were obtained with Matlab, and all subsequent statistical estimation was conducted in R. Note that the present study relies on both Matlab and R, since Matlab is often preferred by the neuroimaging community for preparing data for contribution and fast matrix manipulation, whereas R is preferred for its statistical infrastructure and its popular Shiny web app building framework. The BrainEffex interactive R Shiny app (https://neuroprismlab.shinyapps.io/BrainEffeX/) has been developed to facilitate more detailed visualization, exploration, and downloading of the study-specific effect size maps, meta-analyses, and metadata (DOI: 10.5281/zenodo.16882652; Shearer et al., 2025). The BioImage Suite command line software used for processing is freely available at (https://bioimagesuiteweb.github.io/webapp/index.html).

## Acknowledgements

This work was supported by funding from the National Institute of Mental Health (K99 MH130894; R00 MH130894 to S.N).

Data were provided by numerous sources, and we gratefully acknowledge the efforts of multiple teams in collecting, preparing, and sharing this data. Data were provided in part by the Human Connectome Project, WU-Minn Consortium (Principal Investigators: David Van Essen and Kamil Ugurbil; 1U54MH091657) funded by the 16 NIH Institutes and Centers that support the NIH Blueprint for Neuroscience Research; and by the McDonnell Center for Systems Neuroscience at Washington University. This research was conducted using the UK Biobank Resource under application number 140089. The Philadelphia Neurodevelopmental Cohort data were obtained from the database of Genotypes and Phenotypes (dbGaP) through dbGaP accession number phs000607.v1.p1. Support for the collection of the PNC dataset was provided by grants RC2 MH089983 and RC2 MH089924 from the National Institute of Mental Health, with additional support for genotyping from the Children’s Hospital of Philadelphia. The Healthy Brain Network data were provided by the Child Mind Institute with support from the NIMH (awards to M.P. Milham). The Southwest University Longitudinal Imaging Multimodal (SLIM) dataset was supported by grants from the National Natural Science Foundation of China (31271087, 31470981, 31571137, 31500885). Data used in the preparation of this article were obtained from the Adolescent Brain and Cognitive Development (ABCD) Study (https://abcdstudy.org), held in the NIMH Data Archive (NDA). The ABCD Study is supported by the National Institutes of Health and additional federal partners under award numbers U01DA041048, U01DA050989, U01DA051016, U01DA041022, U01DA051018, U01DA051037, U01DA050987, U01DA041174, U01DA041106, U01DA041117, U01DA041028, U01DA041134, U01DA050988, U01DA051039, U01DA041156, U01DA041025, U01DA041120, U01DA051038, U01DA041148, U01DA041093, U01DA041089, U24DA041123, U24DA041147. A full list of supporters is available at https://abcdstudy.org/federal-partners.html. A listing of participating sites and a complete listing of the study investigators can be found at https://abcdstudy.org/consortium_members/. ABCD consortium investigators provided data but did not participate in the analysis or writing of this report. This manuscript reflects the views of the authors and may not reflect the opinions or views of the NIH or ABCD consortium investigators.

## Contribution statement

S.N. conceived of the study. D.S. and S.N. supervised the study. H.S. and S.N. wrote the code for performing all analyses described in the study and conceived of the data contribution formats, with input from M.R. on testing data contribution and A.F. on visualization. S.N. created the cross-brain effect size estimation procedure with regular input from J.C., A.M., F.C., and T.N., and T.N. derived the procedures for estimating cross-brain power under multiple testing control. S.N. wrote the manuscript with regular input from J.C. and D.S. and assistance from H.S., A.F., and F.C. All other authors contributed subject-level data and performed any relevant preprocessing. All authors provided input on the manuscript.

