## Supplementary Methods for "The Hidden Landscape of Missed Effects in Human Functional Neuroimaging"

#### Datasets

Seven large-scale datasets were included based on the availability of functional neuroimaging data and sample sizes >100 participants to provide precise effect size estimates. Datasets were selected that included functional magnetic resonance imaging data and at least a "large" sample size (>400 subjects; cf. (Horien et al., 2021)) to provide as precise effect size estimates as possible. One additional dataset with fewer subjects (>100 subjects) was included to represent clinical populations explicitly.

Detailed information regarding study characteristics is provided in **Supplementary Table 1** and fMRI tasks and scan durations is provided in **Supplementary Table 2**. In brief, the following data was used from each dataset:

The *Human Connectome Project (HCP) Young Adults* dataset provided task (500 Subjects Release) and resting-state (1200 Subjects Release) fMRI data from 1,200 healthy adults aged 22-35. This dataset was used to estimate task-based activation, contrasts between task and rest, and resting-state connectivity associations with cognitive and demographic variables.

The *Human Connectome Project Early Psychosis (HCP-EP)* dataset (Release 1.1) provided resting-state fMRI data from young adults with early psychosis and matched controls. This dataset was used to estimate clinical group differences in resting-state connectivity.

The *UK Biobank (UKB)* dataset (October 2023 Release) provided resting-state fMRI data from approximately 40,000 participants aged 40-69. This dataset was used to estimate resting-state connectivity associations with cognitive, physical, and demographic measures.

The *Adolescent Brain and Cognitive Development Study (ABCD)* dataset (Release 2.0.1) provided task and resting-state data from participants aged 9-12. This dataset was used to estimate contrasts between task and rest and resting-state connectivity associations with cognitive, psychiatric, physical, and demographic variables.

The *Philadelphia Neurodevelopmental Cohort (PNC)* dataset (January 2014 Release) provided resting-state fMRI from participants aged 8-22. We used resting-state connectivity data and working memory task activation maps.

The *Healthy Brain Network (HBN)* dataset (Releases 1.1 and 2.1) provided resting-state fMRI data from children and adolescents with and without psychiatric diagnoses. We used resting-state connectivity data to examine developmental and clinical associations.

The *Southwest University Longitudinal Imaging Multimodal (SLIM)* dataset (February 2017 Release) provided longitudinal resting-state fMRI data from healthy young adults aged 18-26, contributing to developmental connectivity studies.

These datasets capture a broad range of developmental stages, from childhood and adolescence (PNC, ABCD, HBN) to young adulthood (SLIM, HCP) and mid-to-older adulthood (UKB). They include both typically developing populations (HCP, UKB, ABCD, HBN) and clinical or high-risk cohorts (HCP-EP, PNC, HBN, SLIM). Imaging protocols vary, with most datasets including resting-state fMRI to examine intrinsic functional connectivity. Others, such as HCP and ABCD, include task-based fMRI paradigms spanning cognitive, emotional, and motor domains. Additionally, these datasets differ in their breadth of phenotypic assessments, with UKB and ABCD incorporating extensive demographic, cognitive, and health-related measures, and others, such as PNC, including detailed psychiatric and substance use assessments. This diversity allows for a comprehensive examination of factors influencing brain function across the lifespan and in both normative and clinical populations.

**Studies**

Here, a "study" refers to a particular investigation into a phenomenon of interest resulting in a unique effect size map. For example, a study might examine the correlation of a specific phenotype with resting-state fMRI data, differences in functional connectivity between clinical groups, or differences in functional connectivity during task execution versus rest. As such, each study here can be uniquely identified by dataset, map type, and outcome category. The "effect size database" refers to the complete collection of studies. Studies in the current effect size database were selected to reflect as wide a range of outcome categories as possible. The database is particularly enriched in functional connectivity studies, correlational studies, and psychological (cognitive or psychiatric) outcomes. A detailed list of study characteristics is provided in **Supplementary Table 1**. A brief overview is provided as follows:

*HCP Studies:* The following studies were used from the HCP young adults dataset: For the FC task-based studies, motion task-based activation, gambling task-based activation, relational task-based activation, social task-based activation, and working memory task-based activation were all compared to rest data FC. For the association studies, resting-state FC associations with fluid intelligence (Penn Progressive Matrices) and processing speed.

The following tasks and contrasts were selected (Glasser et al., 2013) were selected to reflect a range of task-based effects, as described in (Noble et al., 2020): the Theory of Mind versus Random contrast from the Social task (SOCIAL COPE 6: TOM-RANDOM; N = 484), the Face versus Other contrast from the Working Memory task (WM COPE 20: FACE-AVG; N = 493), the Relational versus Match contrast from the Relational task (RELATIONAL COPE 4: REL-MATCH; N = 480), the Reward versus Punishment contrast from the Gambling task (GAMBLING COPE 6: REWARD-PUNISH; N = 491), and the Faces versus Shapes contrast from the Emotion task (EMOTION COPE 3: FACES-SHAPES; N = 482).

*PNC Studies:* Studies from the PNC dataset examined resting-state FC associations with psychiatric symptom screening questions across multiple domains: attention deficit disorder, depression, eating disorders, generalized anxiety, mania/hypomania, obsessive-compulsive behaviors, panic disorder, psychosis spectrum symptoms (SIPS-PRIME), and a sex FC difference study.

*SLIM Studies:* The SLIM dataset examined resting-state FC associations with state anxiety measured via the State-Trait Anxiety Inventory (STAI) and sex FC difference.

*HBN Studies:* Studies from the HBN dataset examined resting-state FC associations with the following NIH Toolbox cognitive measures: card sorting, flanker inhibitory control, list sorting with working memory, processing speed, Social Responsiveness Scale scores, and diagnostic categories.

*ABCD Studies:* The ABCD dataset examined resting-state FC associations with age, BMI, CBCL syndrome scales (aggressive, anxious/depressed, attention, externalizing, internalizing, rule-breaking, social, somatic, thought, withdrawn/depressed), NIH Toolbox fluid cognition composite, and WISC-V matrix reasoning. Additional studies examined FC differences between sex and FC contrast between resting-state and task conditions (MID, n-back, SST). Details regarding the task designs are provided in **Supplementary Table 2**.

*UKB Studies:* The UKB neuroimaging sample examined resting-state FC associations with age and fluid intelligence (from both verbal and numerical reasoning tests).

**Subject-level preprocessing & analysis**

Most datasets were preprocessed using the “standard” Yale MRRC in-house pipeline for estimating functional connectivity except the following four map & dataset combinations: (1) activation maps from the HCP dataset, (2) functional connectivity maps from the HCP dataset (partial pipeline), (3) functional connectivity from the SLIM dataset, and (4) functional connectivity from the UKB dataset.

*Yale MRRC Processing and Functional Connectivity Pipeline (ABCD, HBN, and PNC)*

For ABCD, HBN, and PNC, structural images were skull-stripped (FSL), and functional images were motion-corrected and slicetime-corrected (SPM8). Subsequent steps occurred in BioImage Suite (Joshi et al., 2011), including nonlinear registration of structural images to MNI space via iterative group template optimization and linear registration of mean functional images to skull-stripped structural images. The following noise covariates were regressed from the data: linear and quadratic drift, a 24-parameter model of head motion (including six rigid body motion parameters, six temporal derivatives, and all terms squared), mean cerebrospinal fluid, mean white matter, and mean global signal. Data were then temporally smoothed with a Gaussian filter (cutoff frequency ≈ 0.12Hz). Functional connectivity was calculated as Fisher z-transformed Pearson correlations between mean timecourses for each region pair in the Shen 268-node whole-brain atlas (Shen et al., 2013).

*Additional processing details for HCP functional connectivity*

Minimally processed data was obtained from the HCP and subsequently processing with BioImage Suite. HCP minimal preprocessing included gradient distortion correction, motion correction, fieldmap-based distortion correction, and brain-boundary-based registration of functional to structural scans (Glasser et al. 2013). As above, subsequent processing in BioImage Suite included regression of nuisance parameters (linear and quadratic drift, the 24-parameter model of head motion, mean cerebrospinal fluid signal, mean white matter signal, and mean global signal), temporal smoothing (cutoff frequency ≈ 0.12Hz), and nonlinear registration to MNI space. Functional connectivity was calculated as Fisher z-transformed Pearson correlations between Shen 268 regions separately for right-left and left-right phase encoding scans and then averaged.

*Additional processing details for HCP task-based activation*

Subject-level activation beta coefficients were obtained from the HCP repository, which were previously generated by HCP as follows. After minimally processing data as above, additional processing (Barch et al., 2013) included nonlinear registration to MNI152 space (FNIRT), grand-mean intensity normalization, and spatial smoothing (4mm FWHM Gaussian kernel). Activations were estimated via regression with task-specific parameters convolved with a canonical hymodynamic response function. Temporal derivatives of predictors were included as confounds. Both time series and GLM design were highpass filtered (Gaussian-weighted, 200s cutoff) and data were prewhitened.

*Additional processing details for SLIM functional connectivity*

Functional connectivity maps were obtained from the SLIM repository, which were previously generated by SLIM as follows (Liu et al., 2017). The first 10 volumes of the functional images were discarded to account for signal equilibrium and participants' adaptation to their immediate environment. Slice timing, motion correction and spatial normalization to a standard template were applied to the remaining 232 images. Images were then re-sampled to 3mm cubic voxels, followed by spatial smoothing (6mm FWHM). Data were linearly detrended and temporally smoothed using a band pass filter (0.01-0.08 Hz). The following noise covariates were regressed from the data: six head motion parameters, mean cerebrospinal fluid, white matter, and global signal.

*Additional processing details for UKB functional connectivity*

Functional connectivity maps were obtained from the UKB repository, which were previously generated by UKB using the automated UK Biobank pipeline as follows

(Alfaro-Almagro et al., 2018). First, the data was nonlinearly warped to MNI152 space using FNIRT. Next, tissue-type segmentation was applied via FAST to obtain cerebrospinal fluid, grey matter, and white matter boundaries, as well as partial-volume images for each tissue type. Next, the Melodic was used for the following steps: motion correction using MCFLIRT, grand-mean intensity normalisation of the entire 4D dataset by a single multiplicative factor, highpass temporal filtering (Gaussian-weighted least-squares regression with sigma=50.0s), fieldmap-based distortion correction, and gradient distortion correction. ICA+FIX was used to remove structured artifacts. Resting-state data underwent group-ICA (MELODIC) at dimensionality 25 to identify functional network nodes. Non-neural components were removed. Subject-specific network matrices were estimated via dual regression, calculating full correlations between node timecourses (Alfaro-Almagro et al., 2018).

**Mass univariate group-level test statistics**

Subject-level functional connectivity and task-based activation maps were then used for group-level analysis in conjunction with any outcome variables. Each study involved one of three group-level statistical tests, described below. Each test was further adjusted using four motion adjustment strategies and three scales of inference, resulting in 12 statistical maps per study. All tests, adjustments, and levels of inference were selected to reflect common practices in the field, with the exception of the adjustment procedures for the multivariate tests, which were selected to as closely match the univariate tests as possible.

In order to automate group-level analysis here and in future studies, subject-level data was organized in the BrainEffeX contributor format. This format enables the data to be ingested by calculate_effex and subsequent tools for effect size analysis and ultimate inclusion in the Shiny app (for more information about contribution standards, see Shearer et al. (2025).

*Classes of group-level tests*

Each study involved one of three tests: one- or paired-sample t-test (“t”), two-sample t-test (“t2”), and correlation (“r”). A general linear model can be formulated for all tests as follows:

$$Y = X\beta + \epsilon$$

with variables defined according to the test:

*1- or paired-sample t-test:* Let $Y \in R^{n\times 1}$ be a brain variable (i.e., voxel, edge, or network; difference scores are used for the paired condition) measured for $n$ subjects and $X = 1$ be a column vector of ones,

*2-sample t-test:* Let $Y \in R^{n\times 1}$ be a brain variable measured for $n$ subjects and $X \in R^{n\times 2}$ be a design matrix where the first column consists of ones and the second column is a binary indicator variable representing the group assignments for each of the $n$ subjects,

*Correlation:* Let $Y \in R^{n \times 1}$ be a score variable measured for $n$ subjects and $X \in R^{n \times 2}$ be a design matrix where the first column consists of ones and the second column contains a brain variable measured for $n$ subjects.

Then $\beta \in R^1$ (for "t") or $\beta \in R^2$ (for "t2" or "r") are the regression coefficients and $\epsilon \in R^{n \times 1}$ is the residual error. $t$ statistics are calculated for "t" and "t2" tests and $r$ statistics are calculated for "r" tests as described below. Here we introduce notation for univariate tests without deconfounding; we will later extend this notation to the deconfounding and multivariate cases. Notation used in this and the following sections follows conventions established in (Winkler et al., 2014).

*Motion adjustment strategies*

Three strategies were used to explore the effects of correcting motion during group-level inference: no motion correction ("none"), inclusion of motion as a predictor ("motion regression"), and exclusion of subjects with high motion ("thresholding"). Note that these group-level motion adjustment strategies were included in addition to earlier rigid-body motion correction and motion time series regression steps described in the preprocessing section.

*1. None*

The "none" strategy entailed standard calculation of univariate $r$ and $t$ statistics without inclusion of motion as a regressor.

*2. Motion regression (statistical control)*

A partitioned model that includes confounds can be formulated as

$$Y = X\beta + Z\gamma + \epsilon \,,$$

where $Z \in R^{n \times 1}$ represents motion measured by the mean frame-to-frame displacement (mFFD) for $n$ subjects.

For correlation, "statistical control" is represented by the semipartial correlation statistic

$$r_{Y(X \sim Z)} = \frac{r_{YX} - r_{YZ} r_{XZ}}{\sqrt{1 - r_{XZ}^2}} \,.$$

where $r_{YX}$ , $r_{YZ}$ and $r_{XZ}$ are the pairwise correlations between variables. This facilitates the estimation of ${R_{Y(X \sim Z)}}^2$, the proportion of variance in $Y$ that can be explained by $X$ above and beyond the variance explained by $Z$ (i.e., above that explained by motion).

${R_{Y(X\sim Z)}}^2$ is the standard measure of "variance explained" by a predictor that is obtained when fitting a multiple regression model.

For the two-sample t-test, the semipartial point biserial correlation is estimated and converted to a t-statistic as described in (Kim, 2022):

$$t_{Y(X\sim Z)} = r_{Y(X\sim Z)} \frac{\sqrt{n-2-g}}{\sqrt{1-{r_{Y(X\sim Z)}}^2}} = r_{Y(X\sim Z)} \frac{\sqrt{n-3}}{\sqrt{1-{r_{Y(X\sim Z)}}^2}}$$

where $g = 1$ is the number of confounds. Note that the p-value can also be estimated as $p = 2\Phi_t(-|t|, n-2-g) = 2\Phi_t(-|t|, n-3)$, where $\Phi_t(\cdot)$ is the cumulative density function of the Student's t distribution with degrees of freedom $n-3$.

For the one-sample t-test, "statistical control" is represented by the typical t-statistic obtained for the intercept of the full multiple regression model. Namely, using an unpartitioned model

$$Y = X\beta + \epsilon \text{ where } X \leftarrow [X, Z] \text{ and } \beta \leftarrow [\beta, \gamma],$$

the t-statistic is calculated as

$$\beta = (X'X)^{-1}X'Y$$

$$SE(\beta) = \frac{1}{n-p}\sum_{i=1}^{n} (Y_i - \beta X_i)^2 = \frac{1}{n-2}\sum_{i=1}^{n} (Y_i - \beta X_i)^2$$

$$t = \beta / SE(\beta)$$

where $p = 2$ is the number of estimated parameters with a summation over each subject $i$. Like $R^2$, the t-statistic can be used to obtain the amount of variance explained by freely estimating an intercept compared to a null (zero) intercept model.

*3. Thresholding*

For the "thresholding" strategy, a threshold was applied to exclude high motion subjects and then estimation proceeded without further consideration of motion. Two different thresholds were examined. For the first, subjects with mFFD>0.1 mm were discarded, based on the prior observation that 95% of healthy young adult subjects exhibit motion below this level (unpublished).

*Scales of inference*

Finally, for each study, results were additionally examined at three scales of inference: at the level of individual edges or voxels ("element-level"), larger scale collections of brain areas ("network-level"), and across the whole-brain ("whole brain-level").

*1. Element- and network-level test statistics*

For element-level statistics, a test statistic was calculated for each edge or voxel in a "mass univariate" fashion. Specifically, for each collection of $k$ edges or voxels, $k$ separate models were formulated and separately used to calculate $k$ statistics.

*2. Network-level test statistics*

Network-level statistics are estimated as for element-level statistics, except voxels or edges were first averaged within node groups or subnetworks of the Shen atlas respectively prior to calculation of the test statistic. In addition to providing a parcellation of the brain into 268 regions, the Shen atlas also provides an assignment of each region to one of 10 node groups (i.e., large-scale networks). For network-level analysis, Edges were first averaged within subnetworks of the Shen atlas to yield a set of 55 subnetwork measurements (i.e., entries in lower triangle plus diagonal of a 10-"node group" matrix = 10 node groups x (10 node groups + 1) / 2 = 55 subnetworks; cf. (Noble et al., 2017)).

*3. Whole brain-level (multivariate) test statistics*

Multivariate test statistics spanning the whole brain were calculated using similar procedures as described above, except that $Y$ for the 1- and 2-sample t-tests is instead defined as $Y \in R^{n \times k}$ and represents a collection of $k$ voxels or edges measured from $n$ subjects, and $X$ for the correlation test is instead defined as $X \in R^{n \times k}$ and represents a collection of $k$ voxels or edges measured from $n$ subjects.

First, any required motion adjustment was performed prior to any dimensionality reduction or calculation of statistics, as recommended by (Winkler et al., 2020). Mean-centered motion was regressed from variables representing brain data and scores by:

$$Z_c = Z - \bar{Z}$$
$$P = Z_c(Z_c{}'Z_c)^{-1}Z_c{}'$$
$$\tilde{Y} = Y - R_Z Y$$
$$\tilde{X} = X - R_Z X$$

where $Z_c$ is the mean-centered confound vector, $P$ is the prediction matrix, and $\tilde{Y}$ and $\tilde{X}$ are the residualized variables. All motion adjustment strategies were chosen to parallel those used in the univariate case. Specifically, to obtain partial correlation in the "motion regression" case, $Z$ was regressed from both $X$ and $Y$ (cf. "Partial CCA" and "Part CCA" in Table 1 of Winkler et al., 2020). The one case that had to be omitted was the one-sample t-test with "motion regression", since the standard Hotelling T-squared method does not directly accommodate the addition of a nuisance parameter to parallel the procedure used in the univariate "motion regression" case (i.e., $Y \sim 1 + Z$). Finally, for "thresholding", subjects were omitted based on mFFD as above, and for "none", $Z$ was not included in any way.

Next, a reduced representation of the brain data $X$ or $Y$ was calculated in order to allow use of multivariate statistics, with additional reduction to calculate reliable statistics (Helmer et al., 2024). (Helmer et al., 2024) determined empirically that error of an estimated canonical correlation coefficient can be reduced to "reliable" levels by reducing the full set of brain variables to $q = \lfloor n/50 \rfloor$ components via Principal Component Analysis (PCA). As such, the original $X$ or $Y$ is replaced by the top $q$ principle components of a PCA of that variable:

$$\text{for t-tests: } Y^* \leftarrow PCA_q(Y)$$
$$\text{for correlation: } X^* \leftarrow PCA_q(X)$$

where $PCA_q(\cdot)$ represents a principle component analysis operation that returns the top $q$ components, and $Y^* \in R^{n\times q}$ or $X^* \in R^{n\times q}$represents the corresponding score vectors (rows represent the new scores associated with the $q$ components for a single subject). Note that centering is not performed for the one-sample case since the centroid is used for the test.

The multivariate 1-sample t-test statistic was estimated as the Hotelling $t^2$ statistic for the intercept of $Y = \beta_0 + \epsilon$ , taking the square root to obtain $t = \sqrt{t^2}$. This statistic is based on the multivariate distance from the origin and its covariance matrix.

The multivariate statistics for both 2-sample t-test and correlation were estimated using the top canonical variables via Canonical Correlation Analysis. This proceeds by finding the canonical coefficients $A$ and $B$ that maximize the correlation $r$ between

$$u = YA$$
$$v = XB$$

where $u \in R^{n\times 1}$ , and $v \in R^{n\times 1}$ are the the top canonical variables and $r$ is the resulting multivariate test statistic. For the 2-sample t-test, $A \in R^{q\times 1}$ and $B \in R^1$ , whereas for correlation, $A \in R^1$ and $B \in R^{q\times 1}$ .

### Implementation of group-level statistical procedures

*Matlab* was used for group-level inference in order to leverage its innate efficiency with matrix manipulation, as well as because it is a preferred language in the neuroimaging community for preparing data for contribution. Built-in or custom functions were used for all procedures described except as follows. The "Regression_fast" function from the "Fast_Regression" package was modified to expedite coefficient estimation in general and for the mass univariate case. The *T2Hot1* function from the HotellingT2 toolbox (https://www.mathworks.com/matlabcentral/fileexchange/2844-hotellingt2) was modified for the Hotelling T-squared test (Trujillo-Ortiz, 2026).

### Effect size conversion

#### *Effect size measures*

Results were reported as Cohen's $d$ and $R^2$ in order to facilitate interpretation and comparison across study types. Cohen's $d$ reflects standardized differences between groups or from the intercept, and $R^2$ reflects the proportion of variance of the outcome explained by the predictor. In general, Cohen's $d$ and $R^2$ were calculated from the t-statistics ($t$) and correlation coefficients ($r$) as follows, according to the conventions provided in (Cohen, 2013)):

| | | |
|---|---|---|
| 1-sample t-test: $d = \frac{t}{\sqrt{n}}$ | (Cohen 2013, p 72) | N/A |
| 2-sample t-test: $d = t\sqrt{\left(\frac{1}{n_1} + \frac{1}{n_2}\right)}$ | (Cohen 2013, p 67) | $R^2 = \frac{t^2}{t^2+(n-2)}$ |
| Correlation: $d = \frac{2\,r}{\sqrt{(1-r^2)}}$ | (derived from Cohen 2013, p 23) | $R^2 = r^2$ |

with $n$ denoting the sample size of the group for the 1-sample t-test and correlation and $n_1$ and $n_2$denoting the sample size of each group for the 2-sample t-test.

Note that there are multiple estimators of Cohen's $d$ and $R^2$; the above conversions were selected to reflect conventional conversions. Note also that we are using the standard conversion for the correlation coefficient to Cohen's $d$, where d implicitly reflects the change in the outcome associated with two standard deviations change in the predictor (i.e., $\Delta x = 2\sigma$; cf. Mathur & VanderWeele, 2021). We note that this is a typically un-acknowledged assumption underlying a typical convention, but that other choices of $\Delta x$ may also be reasonable and influence the resulting estimated effect size.

Multivariate effect sizes were estimated by treating the canonical correlation or square root of the Hotelling T-squared statistic as a bivariate correlation or t-statistic for subsequent conversion to Cohen's $d$. This compresses the multivariate effect to only the one or two axes used to define the effect and involves an assumption that the sample size would only affect estimates along those axes, but in the absence of another preferred method for estimating and interpreting multivariate effect sizes, this method was chosen as a starting place.

It is also important to note that although all Cohen's $d$ effect sizes share a name, there is an additional barrier between directly comparing Cohen's $d$ from within- and between-sample tests in that the same Cohen's $d$ has different power implications (**SI Fig. 2**).

Despite these different implications, conversion to these effect sizes was chosen as a starting place for comparison and interpretation across studies and may be compared in terms of sample size requirements.

*Uncertainty measures*

Edge-, voxel-, and network-level effect sizes, standard errors (SE), and confidence intervals (CI) were calculated. Confidence intervals (multivariate results only) are estimated using an error threshold $\alpha = 0.05$ ($z = 1.96$), corresponding with 95% control for a variable within a study. Simultaneous confidence intervals are estimated across all $m$ variables using $\alpha = 0.05/m$ in the respective study, corresponding with 95% control across all intervals in the study simultaneously.

For the *one-sample t-test*, Cohen's $d$ standard errors and confidence intervals were estimated after conversion to $d$:

$$SE_d = \sqrt{\frac{1}{n} + \frac{d^2}{2n}}$$

$$t_{crit} = t_{(1 - \alpha/2, df)} \text{ where } df = n - 1$$

$$CI_d = [d - t_{crit} \cdot SE_d, d + t_{crit} \cdot SE_d]$$

where $n$ is the sample size of the group.

For the *two-sample t-test*, Cohen's $d$ and $R^2$ standard errors and confidence intervals were estimated after conversion to $d$ and $R^2$:

$$SE_d = \sqrt{\frac{n_1 + n_2}{n_1 \cdot n_2} + \frac{d^2}{2(n_1 + n_2)}}$$

$$t_{crit} = t_{(1 - \alpha/2, df)} \text{ where } df = n_1 + n_2 - 1$$

$$CI_d = [d - t_{crit} \cdot SE_d, d + t_{crit} \cdot SE_d],$$

$$SE_{R^2} = \sqrt{\frac{4 \cdot R^2 \cdot (1 - R^2)^2 \cdot (n-2)^2}{(n^2 - 1) \cdot (n + 3)}}$$

$$z_{crit} = \Phi^{-1}(1 - \alpha/2)$$

$$CI_{R^2} = [R^2 - z_{crit} \cdot SE_{R^2}, R^2 + z_{crit} \cdot SE_{R^2}]$$

where $n_1$ and $n_2$ are sample size for each group, $n = n_1 + n_2$ is the total sample size, and $\Phi^{-1}$ is the quantile function of the normal distribution.

For *correlation*, Cohen's $d$ and $R^2$ standard errors were estimated after conversion to $d$ and $R^2$, correlation coefficient confidence intervals were estimated directly and then converted to Cohen's $d$ confidence intervals, and $R^2$ confidence intervals were estimated after conversion to $R^2$:

$$SE_d = \sqrt{\frac{n_1 + n_2}{n_1 \cdot n_2} + \frac{d^2}{2(n_1 + n_2)}}$$

$$z_{crit} = \phi^{-1}(1 - \alpha/2)$$

$$CI_r = [tanh\,(tanh^{-1}\,(r) - \frac{z_{crit}}{\sqrt{n-3}}), tanh\,(tanh^{-1}\,(r) + \frac{z_{crit}}{\sqrt{n-3}})]$$

$$CI_d = [\frac{\sigma_X \cdot CI_{r,lower}}{\sqrt{(1 - CI_{r,lower}{}^2)}}, \frac{\sigma_X \cdot CI_{r,upper}}{\sqrt{(1 - CI_{r,upper}{}^2)}}],$$

$$SE_{R^2} = \sqrt{\frac{4 \cdot R^2 \cdot (1 - R^2)^2 \cdot (n - 2)^2}{((n^2 - 1) \cdot (n + 3))}}$$

$$z_{crit} = \phi^{-1}(1 - \alpha/2)$$

$$CI_{R^2} = [R^2 - z_{crit} \cdot SE_{R^2}, R^2 + z_{crit} \cdot SE_{R^2}]$$

where $\sigma_X = 2$ , as specified in the effect size conversion, and $n$ is the sample size.

For reference, a confidence interval has the property that, over many repeated instances of the entire data, there is a 95% probability across samples of containing the true value. Similarly, the simultaneous confidence interval for multiple estimates is the collection of intervals, of many repeated instances, which jointly contain all true values across 95% of samples. Confidence intervals do not directly indicate information about the uncertainty of the point estimates. Simultaneous confidence intervals can be used as one way of providing a "corrected" measure of uncertainty that represents the range of values expected to encompass the true effect for all edges or voxels across an expected proportion of repetitions of an experiment—in short, the more variables, the wider the intervals. When there are many effects being estimated (e.g., one per voxel), it is helpful to control the joint probability of all intervals containing the true value in the same way that it is helpful to control the joint probability of false positives across multiple tests (i.e., multiple comparison correction).

While simultaneous confidence intervals are helpful for visualizing the joint estimate of uncertainty for an individual study, it is currently unclear how they can be used to provide a more precise estimate of the distribution of effect sizes across variables than that given by the point estimates, which, as discussed below, produces inflated magnitudes (see *Methods: Estimating population effect size distribution*).

#### Grouping studies by outcome category

Studies were manually assigned different categories based on the outcome of interest. The “psychological” category included between-subject studies of psychiatric and cognitive outcomes. This category contained psychiatric and cognitive sub-categories. Psychiatric outcomes included all assessments that are currently used in the assistance of diagnosis, whereas cognitive outcomes included all psychological measures of cognition. The “task-based” contrasts formed a second category, where “task-based” is a term of art in fMRI research that typically refers to data collected from subjects while they perform tasks in the scanner. All other variables included those typically associated with measures of the body and were included within the “physical” (i.e., anthropometric) overarching category. This category contained sex/gender, age, and biometric (body mass index) subcategories. Notably, sex and gender were grouped together. Studies frequently measure sex as sex-assigned-at-birth and gender as self-reported gender, and while these are combined here in the interest of grouping related measures, we strongly emphasize that self-reported gender is a social construct and can differ from biological sex.

#### Estimating the true distribution of effect sizes across the brain

*Formulating the problem*

Our main objective is to estimate the distribution of effect sizes across multiple brain areas, which we term the "cross-brain effect size distribution". Here we formulate this objective, and we also illustrate how the common strategy of reporting observed mass univariate effect sizes produces distributions that are predictably wider than the true distribution due to sampling error.

In a given study, the observed effect size $\widehat{\theta_i}$ for a brain variable $i$ (here, a voxel, edge, or network) measured with sampling error is specified using a two-level model, following the notation of Harrer et al. (2021), Chapter 4.1.2:

$$\text{Level 1: } \widehat{\theta_i} = \theta_i + \epsilon_i$$

$$\text{Level 2: } \theta_i = \mu + \eta_i$$

where $\epsilon_i$ (level 1) is the deviation in brain variable $i$ from the true effect size $\theta_i$ due to sampling error, and $\eta_i$ (level 2) is the "true" (i.e., noise-free) deviation of brain variable $i$ from the mean across all $k$ variables across the brain $\mu$ .

Substituting the Level 2 expression in Level 1 produces:

$$\hat{\theta}_i = \mu + \eta_i + \epsilon_i \ .$$

The main objective is to estimate the **cross-brain distribution of effect sizes**, which, here, represents the distribution of effect sizes across brain areas. Assuming normality, this distribution is given by

$$\theta_i \sim \mathrm{N}(\mu, {\tau_\eta}^2)$$

where

$${\tau_\eta}^2 = var(\eta_i)\ .$$

Note how the distribution of $\theta_i$ differs from $\hat{\theta}_i$ , the observed mass univariate effect size distribution often provided in published studies, with the key difference being the inclusion of sampling error in the latter. Namely, the variance of $\hat{\theta}_i$ is greater than the variance of $\theta_i$ , the latter of which being our actual target:

$$var(\hat{\theta}_i) = var(\eta_i + \epsilon_i) > \ var(\eta_i)\ .$$

*Estimation approach*

We have described the problem for a single study, but we can also add a third level representing study-specific deviations consistent with a standard meta-analysis. Then, one would ordinarily estimate the heterogeneity (${\tau_{(2)}}^2$) using multi-level meta-analysis estimation procedures implemented in the *metafor* R package (Veichtbauer, https://wviechtb.github.io/metafor/reference/rma.mv.html#references-1). However, two issues prevent their application here. First, multivariate meta-analysis requires the provision of the sample covariance matrix across brain variables, which is expected to be highly unstable for this application given the large dimensionality of the neuroimaging data. Second, and more importantly, we operate under the strong expectation that even the most similar studies included here are sufficiently different as to exhibit markedly distinct patterns of effects across brain areas, and thus it does not make sense to pool across studies within a brain area.

However, we also anticipate that despite these differences in the marginal effects, the overall shape of the cross-brain effect size distribution will be similar across studies of a similar category. Thus, we introduce an alternative strategy for estimating this distribution, with the intention of obtaining a more global view of the magnitude of effects.

In a given study, assuming that levels 1 and 2 deviations are independent, we can write the observed cross-brain variance as the sum of the true cross-brain variance and the added sampling error:

$$var(\hat{\theta}_i) = var(\eta_i + \epsilon_i) = var(\eta_i) + var(\epsilon_i) \,.$$

In the case of a one-sample t-test, the sampling error is the following function of population variance $\phi^2$ (Casella & Berger, 2002, Statistical Inference)

$$var_{1-sample}(\epsilon_i) = \phi^2/n$$

For the two-sample t-test, the sampling error is a multiple of the above when assuming equal variances (i.e., ${\phi_1}^2 = {\phi_2}^2 = \phi^2$) and sample sizes (i.e., $n_1 = n_2 = n_{total}/2$) across groups:

$$var_{2-sample}(\epsilon_i) = \frac{{\phi_1}^2}{n_1} + \frac{{\phi_2}^2}{n_2} = \frac{2\phi^2}{n_{total}/2} = 4\phi^2/n_{total} \,.$$

Note that correlation effect sizes are converted to two-sample effect sizes for the present study and thus also use $var_{2-sample}(\epsilon_i)$.

Thus, the overall model for observed cross-brain variance is

$$var(\hat{\theta}_i) = {\tau_\eta}^2 + k^2\phi^2/n$$

where $k$ is the number of groups for the test.

In the previous sections we have only been discussing models for a single study. We now use multiple studies to estimate $var(\eta_i)$ and $\phi^2$ as the intercept and slope of the model

$$var(\hat{\theta}_i) \sim 1 + \frac{k^2}{n} \,.$$

Specifically, we can estimate parameters for this model by providing for each study $var(\hat{\theta}_i)$, the sample variance across brain variables in that study, and $\frac{k^2}{n}$ , involving the number of groups for the test and sample size for the study. Note that we now only provide one observation per study (i.e., the sample variance across brain variables for that study) instead of multiple observations (i.e., all brain variables).

We additionally aim to provide tailored estimates of $var(\eta_i)$ for each outcome category, nest observations within datasets, and finally weight the observations by sample size during model fitting. We used meta-regression via the *rma.mv* function in *metafor* to accomplish all of these objectives. Specifically, we allow the intercept to vary by category and dataset (i.e., random intercept model):

$$1 \mid category, dataset * category\ .$$

We also use a fixed slope to maximize power to estimate the slope even in categories with fewer studies. As discussed above, one-sample and two-sample t-tests necessarily have different slopes, and given the few one-sample t-test studies present, we expect the one-sample estimate of $\phi^2$ to be biased towards the two-sample estimate $4\phi^2$, which, in turn, may produce slightly deflated intercept estimates; however, inspection of the fitted model for task connectivity suggests relatively minimal bias in the estimated intercepts. Additionally, as in the *Meta-analysis of spatial data* section, we use the default "nlminb" optimizer and alternative optimizers in the case of lack of convergence in the following order: "Nelder-Mead", "BFGS", "bobyqa", "nloptr", "nlm", "hjk". Finally, similar to the procedure in *Meta-analysis of spatial data*, the standard error of $var(\hat{\theta}_i)$ for each study was estimated to provide the weights for the meta-regression. Specifically, after estimating $var(\hat{\theta}_i)$ for a given study, the standard error of that estimated value was calculated as described in the *Confidence intervals* section and then squared: $var(var(\hat{\theta}_i)) = SE(var(\hat{\theta}_i))^2$.

*Counting effects above a target magnitude*

Given $\theta_i \sim \mathrm{N}(\mu, \tau^2)$, we can estimate both the magnitude of effects and the number of effects above a certain magnitude. In the following we show how we can directly estimate the magnitude of effects assuming $\mu = 0$, although we also used an iterative procedure to obtain the number of effects above a certain magnitude and found them to be approximately equal.

Assuming $\mu = 0$, the number of effects expected above a certain magnitude $\bar{\Theta}$ is:

$$n_{|\theta_i|>\bar{\Theta}} \simeq 2 \cdot k \cdot F_{\mathrm{N}}(-\bar{\Theta})$$

where $k$ is the number of variables, $n_{|\theta_i|>\bar{\Theta}}$ is the number of effects above magnitude $\Theta$, $F_{\mathrm{N}}$ is the CDF of the normal distribution $\mathrm{N}(0, \tau^2)$ and the factor 2 results from symmetry. Furthermore, if $\theta_i$ is a normal distribution, standard reference points can be used: 68% of values fall within one standard deviation $\tau$ of the mean, and thus we report this as the magnitude that the "majority of effects" fall under. Further, 27.2% of values fall within one and two standard deviations, so we report that "about a quarter of effects" fall between $\tau$ and $2\tau$ in magnitude. A "minority" (5%) of effects are expected to surpass $2\tau$.

Finally, the expected true effect size magnitude is derived from the fact that the absolute value of a zero centered normal is a half normal

$$E[|\theta_i|] = \tau\sqrt{\frac{2}{\pi}} \qquad .$$

40% of studies (i.e., 25 studies) were found to exhibit non-normality via Shapiro-Wilks test (p<0.05; based on selecting either a maximum of 5,000 variables from all variables sorted by size and equally spaced, or the full set of variables if less than 5,000). However, it is generally appreciated that normality tests are sensitive to even very trivial departures from normality at large sample sizes. Inspection of quantile-quantile plots for each study suggests that the data exhibit sufficient normality to continue with the normality assumption in the present work, although this assumption should be considered in future research alongside other refinements of the shape and nature of the present effect size model. To aid in future investigation, an alternative estimation procedure is provided below that does not assume normality although it does include other assumptions.

*Relaxing the normality constraint*

We can also derive empirical expressions for the above distributions, without assuming normality in the case of sufficiently large $n$. If we assume $n$ is large enough such that the individual error of each effect size is low enough such that $\widehat{\theta_i}$ has the same shape as $\theta_i$, one has:

$$\theta_i \sim \widehat{\theta_i} \cdot \sqrt{\frac{{\sigma_E}^2}{{\sigma_E}^2 + {\sigma_e}^2/n}}$$

In this case, the real empirical effect size distribution can be estimated by scaling each observation in the empirical distribution $\widehat{\theta_i}$, for each $\theta_i$ in the measured empirical effect size distribution.

$$\theta_i = \widehat{\theta_i} \cdot \sqrt{\frac{{\sigma_E}^2}{{\sigma_E}^2 + {\sigma_e}^2/n}}$$

The true effect empirical cumulative distribution can be then calculated as:

$$F_{\theta_i} = \frac{1}{n} {\Sigma_i}^k I_{\{\bar{\Theta} \geq \theta_i\}}(x)$$

Where $I_{\{\bar{\Theta} \geq \theta_i\}}$ is the indicator function that returns 1 if $\bar{\Theta}$ is greater than $\theta_i$ and zero otherwise.

With this assumption, the number of effects expected above a certain magnitude $\bar{\Theta}$ becomes:

$$n_{|\theta_i|>\bar{\Theta}} = k \cdot ((1 - F_{\theta_i}(\bar{\Theta})) + F_{\theta_i}(-\bar{\Theta}))$$

Where $F_{\theta_i}$ is the CDF of the empirical distribution $\theta_i(0, \tau^2)$.

The expected effect size magnitude can be empirically estimated as:

$$E[|\theta_i|] = \frac{1}{k} {\Sigma_i}^k |\theta_i|$$

The assumptions above assume that the estimator error preserves the shape of the distribution, which is valid for larger population sizes as the estimation error decreases as a function of $n$. However, an alternative parametric assumption lies in assuming a shape for the true empirical effect size distribution.

#### Estimating cross-brain distribution power with multiple testing correction

*1. Notation*

For a single test, with effect size $d$ (Cohen's d), sample size $n$, and significance level $\alpha$, the noncentrality parameter is given by $\delta = d\sqrt{n}$. The power for a one-sided z-test is given by

$$Power = \widetilde{\Phi}\left(\widetilde{\Phi}^{-1}(\alpha) - \delta\right),$$

where $\widetilde{\Phi}(t) = 1 - \Phi(t)$ is the complementary CDF of the standard normal distribution, and $\widetilde{\Phi}^{-1}(\alpha)$ is the critical value for level $\alpha$ test.

For a one-sided t-test with $n - 1$ degrees of freedom, the power is given by

$$Power = \tilde{F}_{T(n-1)}\left(\tilde{F}_{T(n-1)}^{-1}(\alpha) - \delta\right),$$

where $\tilde{F}_{T(n-1)}$ is the complementary CDF of the t-distribution with $n - 1$ degrees of freedom, and $\tilde{F}_{T(n-1)}^{-1}(\alpha)$ is the critical value for level $\alpha$ test.

*2. Multiple Testing Testing Set Up*

For $m$ tests, $m_0$ true null hypotheses, and $m_1 = m - m_0$ true alternative hypotheses, we define:

- $H_j$ as the null hypothesis indicator for test $j$, $j = 1, \dots, m$, $H_j = 0$ if null is true $H_j = 1$ if there is a signal present,
- $p_j$ as the p-value for test $j$,
- $p'$ be the p-value threshold for rejecting null hypotheses.

Note that $p'$ may be fixed (e.g. Bonferroni correction) or data-adaptive (e.g. Benjamini-Hochberg FDR), i.e. rejecting null hypothesis $H_j$ if $p_j \leq p'$. (We specifically avoid using $\alpha$ here to avoid confusion with the per-test significance level, or a desired level of FDR control.)

There are multiple measures of power in the multiple testing setting, e.g. power of detecting at least one true effect ('familywise power'), power of detecting at least $K$ effects, power of detecting some proportion of true effects, etc. However, for simplicity, we will just focus on the average power and proportion of adequately powered tests.

*2.1 Average Power - Constant Effect*

Average power is the expected true positive rate (TPR), i.e., the expected proportion of rejected nulls among the tests where the alternative is true:

$$Average\ Power = E\left[\frac{\sum_{j=1}^{m} H_j I(p_j \le p')}{m_1}\right].$$

If all the effects sizes are identical, e.g. $d$ for a one-sample Z-test, the average power is equal to the classical power for a single test with noncentrality parameter $\delta = d\sqrt{n}$,

$$\tilde{\Phi}\left(\tilde{\Phi}^{-1}(p') - \delta\right).$$

*2.2 Average Power - Distribution of Effects*

We now consider the case where the effect sizes $d$ vary according to some distribution with CDF $F_D$. Then the average power is given by integrating over the distribution of effect sizes; for the one-sample one-sided case:

$$Average\ Power = \int \tilde{\Phi}\left(\tilde{\Phi}^{-1}(p') - d\sqrt{n}\right) dF_D(d).$$

For the particular case where the true one-sample effect sizes are normally distributed with mean $\mu_d$ and variance $\sigma_d^2$, this expression is the convolution of two normal CDFs and has the closed form

$$Average\ Power = \tilde{\Phi}\left(\frac{\tilde{\Phi}^{-1}(p') - \mu_d\sqrt{n}}{\sqrt{1 + n\sigma_{d\delta}^2}}\right).$$

*2.3 Proportion of Adequately Powered Tests - Distribution of Effects*

Our goal is to determine the proportion of effects that have power >=80% at a given sample size $n^*$ and an error level $p'$. We'll start by writing the value of the specific $\delta^*$ that can be detected at a significance level $p'$ by rearranging the first equation of this section (see *1. Notation*):

$$\delta^* = \tilde{\Phi}^{-1}(p') - \Phi^{-1}(1 - Power)$$

Recall that the Type II Error Rate $\beta = 1 - Power$ to obtain:

$$\delta^* = \tilde{\Phi}^{-1}(p') - \Phi^{-1}(\beta)$$

We now turn our attention to a distribution of effect sizes $\delta \sim N\left(\mu_d\sqrt{n}, n\sigma_d^2\right)$. We determine the proportion of effects that have power above a target level as the proportion of the distribution $\delta$ that surpasses the critical effect size value $\delta^*$ detectable at the specified power level. We integrate the PDF of $\delta$ from $\delta^*$ to inf, taking $F_\delta$ as the CDF of $\delta$ , then standardize $\delta^*$ to derive an expression in terms of $\Phi$ :

$$P(\delta > \delta^*) = 1 - F_\delta(\delta^*)$$
$$= 1 - \Phi\left(\frac{\delta^* - \mu_d\sqrt{n}}{\sqrt{n}\sigma_d}\right)$$
$$P(\delta > \delta^*) = \tilde{\Phi}\left(\frac{\tilde{\Phi}^{-1}(p') - \Phi^{-1}(\beta) - \mu_d\sqrt{n}}{\sqrt{n}\sigma_d}\right).$$

*2.4 Extension to 2-Sided and 2-Sample Tests*

*2.4.1 2-Sample, 1-Sided Tests*

In the case of a 2-sample test, the noncentrality parameter is $\delta = \left(\frac{1}{n_1} + \frac{1}{n_2}\right)^{-\frac{1}{2}} d$ with distribution $\delta \sim N\left(\left(\frac{1}{n_1} + \frac{1}{n_2}\right)^{-\frac{1}{2}} \mu_d, \left(\frac{1}{n_1} + \frac{1}{n_2}\right)^{-1} \sigma_d^2\right)$, leading to

$$Average\ Power = \widetilde{\Phi}\left(\frac{\widetilde{\Phi}^{-1}(p') - \left(\frac{1}{n_1} + \frac{1}{n_2}\right)^{-\frac{1}{2}} \mu_d}{\sqrt{1 + \left(\frac{1}{n_1} + \frac{1}{n_2}\right)^{-1} \sigma_d^2}}\right)$$

and

$$P(\delta > \delta^*) = \widetilde{\Phi}\left(\frac{\left(\widetilde{\Phi}^{-1}(p') - \Phi^{-1}(\beta)\right) - \left(\frac{1}{n_1} + \frac{1}{n_2}\right)^{-\frac{1}{2}} \mu_d}{\left(\frac{1}{n_1} + \frac{1}{n_2}\right)^{-\frac{1}{2}} \sigma_d}\right).$$

If we assume equal sample sizes ($n_1 = n_2 = n_{total}/2$), then

$$\delta \sim N\left(\sqrt{\frac{n_{total}}{4}} \mu_d, \frac{n_{total}}{4} \sigma_d^2\right),$$

$$Average\ Power = \widetilde{\Phi}\left(\frac{\widetilde{\Phi}^{-1}(p') - \sqrt{\frac{n_{total}}{4}} \mu_d}{\sqrt{1 + \frac{n_{total}}{4} \sigma_d^2}}\right),$$

and

$$P(\delta > \delta^*) = \widetilde{\Phi}\left(\frac{\left(\widetilde{\Phi}^{-1}(p') - \Phi^{-1}(\beta)\right) - \sqrt{\frac{n_{total}}{4}} \mu_d}{\sqrt{\frac{n_{total}}{4}} \sigma_d}\right).$$

Note that this will yield the same result for Average Power and $P(\delta > \delta^*)$ as using the expressions for a one-sample test with $n = \frac{n_{total}}{4}$.

*2.4.2 1-Sample, 2-Sided Tests*

For a 2-sided z-test, the Type I Error Rate is defined over both directions:

$$\alpha = P(|Z| > z_{critical}) = 2P(Z > z_{critical}),$$

where $z_{critical}$ is positive and the rightmost expression is due to symmetry,$P(Z > z_{critical}) = P(Z < z_{critical}) = \frac{1}{2}P(|Z| > z_{critical})$, giving the critical (significance) threshold

$$z_{critical} = \widetilde{\Phi}^{-1}(p'/2).$$

For a true effect, we take the probability of observing a statistic with magnitude greater that $z_{critical}$ (either direction) to determine power:

$$Power = \widetilde{\Phi}\big(\widetilde{\Phi}^{-1}(p'/2) - \delta\big) + \Phi\big(-\widetilde{\Phi}^{-1}(p'/2) - \delta\big).$$

Now considering a distribution of effect sizes $\delta \sim N\big(\mu_d\sqrt{n}, n\sigma_d^2\big)$ , we determine average power by integrating over the distribution of d as before:

$$Average\ Power = \int \big[\widetilde{\Phi}\big(\widetilde{\Phi}^{-1}(p'/2) - d\sqrt{n}\big) + \Phi\big(-\widetilde{\Phi}^{-1}(p'/2) - d\sqrt{n}\big)\big]dF_D(d)$$

$$= \widetilde{\Phi}\left(\frac{\widetilde{\Phi}^{-1}(p'/2) - \mu_d\sqrt{n}}{\sqrt{1 + n\sigma_d^2}}\right) + \Phi\left(\frac{-\widetilde{\Phi}^{-1}(p'/2) - \mu_d\sqrt{n}}{\sqrt{1 + n\sigma_d^2}}\right).$$

For $\mu_d \approx 0$, this simplifies to the following by symmetry

$$Average\ Power = 2\widetilde{\Phi}\left(\frac{\widetilde{\Phi}^{-1}(p'/2) - \mu_d\sqrt{n}}{\sqrt{1 + n\sigma_d^2}}\right).$$

To find the proportion of adequately powered effects, we find the positive noncentrality parameter $\delta^*$ that corresponds with a given power level using Eqn 3 of this section (note that due to symmetry there will also be an equivalent negative noncentrality parameter with the same power level). This is the value of $\delta^*$ that satisfies

$$\delta^* \leftarrow Power = \widetilde{\Phi}\big(\widetilde{\Phi}^{-1}(p'/2) - \delta^*\big) + \Phi(\Phi^{-1}(p'/2) - \delta^*)$$

which can be solved numerically, but here we assume that any counter-tail in the opposite direction of the effect sign is so small as to be approximately zero, thus allowing us to set the lefthand side of the expression to zero and solve for $\delta^*$ as:

$$\delta^* = \widetilde{\Phi}^{-1}(p'/2) - \Phi^{-1}(\beta).$$

Note that this assumption pertains to the univariate sampling distribution of the alternative hypothesis centered at non-zero $\delta^*$, unlike the previous assumption which pertains to the cross-brain distribution $\delta$ centered at 0.

Then, as before, we determine the proportion above $\delta^*$ and below -$\delta^*$ as

$$P(|\delta| > \delta^*) = \widetilde{\Phi}\left(\frac{\delta^* - \mu_d\sqrt{n}}{\sqrt{n}\sigma_d}\right) + \Phi\left(\frac{-\delta^* - \mu_d\sqrt{n}}{\sqrt{n}\sigma_d}\right).$$

If $\mu_d \approx 0$, this again simplifies by symmetry to

$$P(|\delta| > \delta^*) = 2\widetilde{\Phi}\left(\frac{\delta^* - \mu_d\sqrt{n}}{\sqrt{n}\sigma_d}\right),$$

and if we assume the counter-tail of the cross-brain distribution is small enough to be set to zero as above, then

$$P(|\delta| > \delta^*) = 2\widetilde{\Phi}\left(\frac{\widetilde{\Phi}^{-1}(p'/2) - \Phi^{-1}(\beta) - \mu_d\sqrt{n}}{\sqrt{n}\sigma_d}\right).$$

*2.4.3 2-Sample, 2-Sided Tests*

Combining the above sections, we have

$$Average\ Power = 2\widetilde{\Phi}\left(\frac{\widetilde{\Phi}^{-1}(p'/2) - \left(\frac{1}{n_1} + \frac{1}{n_2}\right)^{-\frac{1}{2}}\mu_d}{\sqrt{1 + \left(\frac{1}{n_1} + \frac{1}{n_2}\right)^{-1}\sigma_d^2}}\right)$$

and

$$P(\delta > \delta^*) = 2\widetilde{\Phi}\left(\frac{\left(\widetilde{\Phi}^{-1}(p'/2) - \Phi^{-1}(\beta)\right) - \left(\frac{1}{n_1} + \frac{1}{n_2}\right)^{-\frac{1}{2}}\mu_d}{\left(\frac{1}{n_1} + \frac{1}{n_2}\right)^{-\frac{1}{2}}\sigma_d}\right).$$

*3. Setting the p-value threshold*

For an uncorrected test we'd typically set $p' = \alpha = 0.05$, or for a Bonferroni correction $p' = \alpha_{FWE}/m = 0.05/m$. For BH-FDR, however, $p'$ is random and depends on the data and we need a different approach.

*4. Power for False Discovery Rate*

Following (Genovese and Wasserman, 2002), we conceive of the p-values as being drawn from a mixture of distributions, with proportion $\pi_0$ coming from a uniform distribution, and proportion $\pi_1$ coming from an alternate p-value distribution $F_1$. Using this framework, for large $m$, the FDR threshold $p'$ must satisfy

$$\pi_0 p' + \pi_1 F_1(p') = \frac{p'}{\alpha_{FDR}};$$

note that this doesn't depend on $m$, only on the proportions $\pi_0$ and $\pi_1 = 1 - \pi_0$. This expression is implicit in $p'$ (cannot be solved for $p'$) and for a given $\pi_0$, $\pi_1$, $\alpha_{FDR}$ and effect size reflected in $F_1$, the $p'$ that satisfies the expression must be found numerically.

For a one-sample, one-sided test, $F_1$ p-value distribution is given by

$$F_1(t) = \widetilde{\Phi}\left(\widetilde{\Phi}^{-1}(t) - d\sqrt{n}\right).$$

If there is a distribution of effect sizes, then $F_1$ reflects a convolution of the sampling distribution of the test statistics with this distribution of effect sizes. Like the average power for heterogeneous effects above, if the effect sizes are normally distributed with mean $\mu_d$ and standard deviation $\sigma_d$, then the alternate p-value distribution is given by

$$F_1(t) = \widetilde{\Phi}\left(\frac{\widetilde{\Phi}^{-1}(t) - \mu_d\sqrt{n}}{\sqrt{1 + n\sigma_\delta^2}}\right).$$

*4.1. Extension to 2-Sided and 2-Sample Tests*

*4.1.1 2-Sample, 1-Sided Tests*

Following the logic in 2.4.1, we obtain the following expression for $F_1$ :

$$F_1(t) = \widetilde{\Phi}\left(\frac{\widetilde{\Phi}^{-1}(t) - \left(\frac{1}{n_1} + \frac{1}{n_2}\right)^{-\frac{1}{2}}\mu_d}{\sqrt{1 + \left(\frac{1}{n_1} + \frac{1}{n_2}\right)^{-1}\sigma_d^2}}\right).$$

*4.1.2 1-Sample, 2-Sided Tests*

We start with the extension to a 2-sided z-test under a fixed positive effect. Let $Z \sim N(\delta, 1)$ under $H_1$ with $\delta > 0$; the two-sided p-value is

$$P = 2\{1 - \Phi(|Z|)\} = 2\Phi(-|Z|).$$

For a given $u \in (0,1)$, the non-null p-value CDF is

$$F_1(u) = Pr(P \leq t \mid H_1).$$

Two-sided p-value $\leq t$ is equivalent to $|Z|$ exceeding the critical value $u$:

$$P \leq t \quad \Leftrightarrow \quad 2\{1 - \Phi(|Z|)\} \leq t \quad \Leftrightarrow \quad |Z| \geq \widetilde{\Phi}^{-1}(t/2).$$

Thus

$$F_1(t) = Pr\left(|Z| \geq \widetilde{\Phi}^{-1}(t/2) \mid Z \sim N(\delta, 1)\right).$$

Breaking this into the two tails gives the distribution

$$F_1(t) = Pr\left(Z \geq \widetilde{\Phi}^{-1}(t/2)\right) + Pr\left(Z \leq -\widetilde{\Phi}^{-1}(t/2)\right)$$
$$= \widetilde{\Phi}\left(\widetilde{\Phi}^{-1}(t/2) - \delta\right) + \Phi\left(-\widetilde{\Phi}^{-1}(t/2) - \delta\right).$$

For $\delta \approx 0$ we have one approximation

$$F_1(u) \approx 2\widetilde{\Phi}\left(\widetilde{\Phi}^{-1}(t/2) - \delta\right)$$

but a more sensible approximation is for $\delta > 0$, and we neglect the lower tail contribution

$$F_1(u) \approx \widetilde{\Phi}\left(\widetilde{\Phi}^{-1}(t/2) - \delta\right).$$

Following the logic in 2.4.2, we extend this expression to model a distribution of effects as:

$$F_1(t) = 2\widetilde{\Phi}\left(\frac{\widetilde{\Phi}^{-1}(t/2) - \mu_d\sqrt{n}}{\sqrt{1 + n\sigma_d^2}}\right).$$

*4.1.3 2-Sample, 2-Sided Tests*

Following the logic in 4.1.1 and 4.1.2, we obtain the following expression for $F_1$:

$$F_1(t) = 2\widetilde{\Phi}\left(\frac{\widetilde{\Phi}^{-1}(t/2) - \left(\frac{1}{n_1} + \frac{1}{n_2}\right)^{-\frac{1}{2}} \mu_d}{\sqrt{1 + \left(\frac{1}{n_1} + \frac{1}{n_2}\right)^{-1} \sigma_d^2}}\right).$$

*5. Implementation & Demonstration*

Procedures for power estimation over a distribution of effects with multiple testing correction have been implemented in the *crossbrain_effects* repository (https://github.com/neuroprismlab/crossbrain_effects). Two illustrative cases are provided below.

First, we examine the influence of different proportions of null signals on power with homogeneous effect sizes (**SI Fig. 7**). This nicely shows how FDR has much higher power than Bonferroni correction, especially when there are more non-null signals (lower $\pi_0$). FDR is not as powerful as uncorrected testing, but the difference is not large when there are many non-null signals.

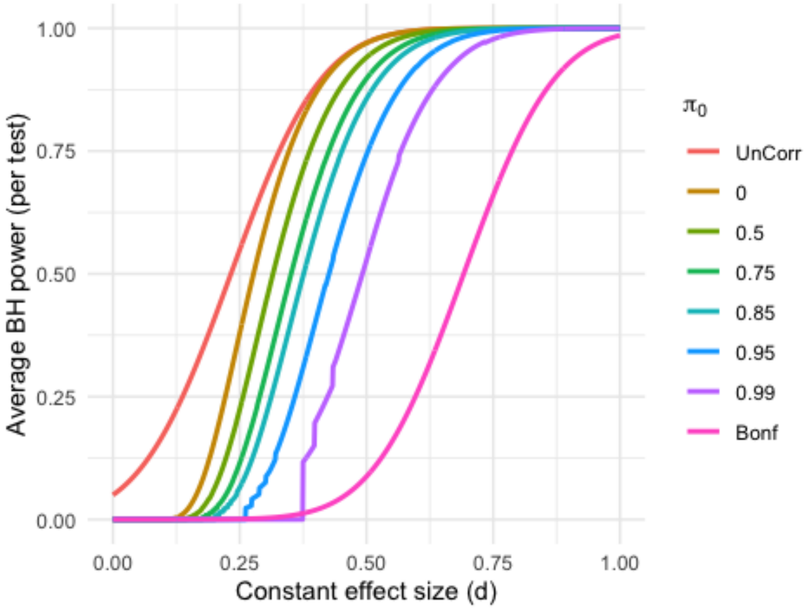


**SI Fig. 7.** Power curves across $\pi_0$ values (proportion null signal), assuming constant effect size for all non-null signals. Sample size = 50, $\alpha_{FDR}$ = 0.05.

Second, we examine the influence of signal heterogeneity on power in the case of heterogeneous effect sizes (**SI Fig. 8**). This shows the flattening effect that signal heterogeneity has on the power curve. But recall the crucial difference in interpretation: While the previous homogeneous plot is for the case that all signal voxels share the exact same effect size $d$, here there is a distribution of effect sizes and we are plotting against the mean effect size about which the actual effect sizes are distributed.

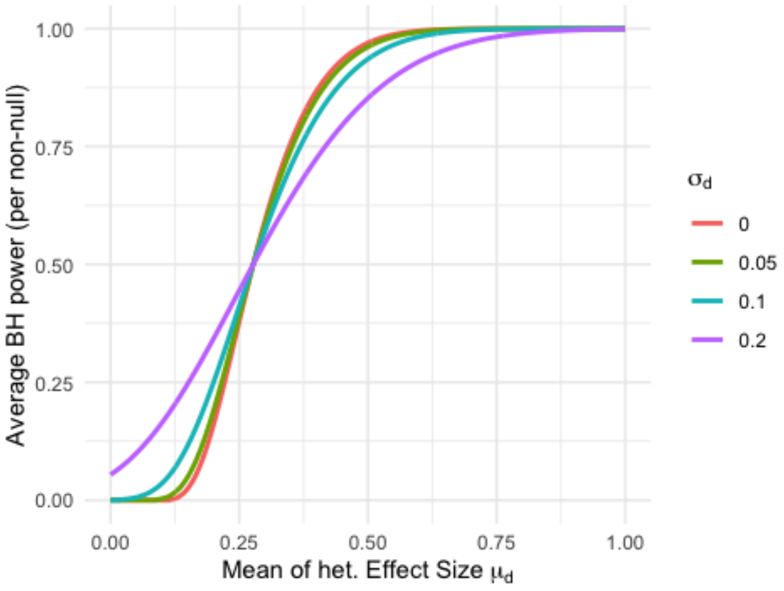

**SI Fig. 8.** Power curves for $\pi_0 = 0$ (true signal everywhere) across different signal heterogeneity values. Sample size = 50, $\alpha_{FDR}$ = 0.05.

### Errors from study planning based on uncorrected effect sizes

*Simulated studies based on corrected and uncorrected effect sizes*

For a given outcome category, ground truth effect sizes for 500 brain regions were drawn from $N(0, {\tau_\eta}^2)$ (level 2) and subject-specific error for 10,000 subjects was drawn from $N(0, \phi^2)$ (level 1), yielding a master population with observed effect sizes for each region.

The following steps were conducted for four study planning strategies ("basis sample size," "basis max effect," "basis average effect," and "corrected effect") at each of six *a priori* sample sizes $n = [25, 50, 100, 500, 1{,}000, 5{,}000]$.

For strategies using basis studies, a basis study was conducted by drawing the *a priori* sample size from the population and performing a one-sided, one-sample t-test. The main study sample size and researcher-expected true positives ($TP_E$) were defined per strategy as follows. For the *"optimistic literature"* strategy, sample size was calculated to obtain adequate power for the maximum significant effect via conventional power analysis (i.e., pwr.t.test), and $TP_E = 1 \times \beta$, where $\beta$ is target power. For *"conservative literature,"* sample size was calculated to obtain adequate power for the minimum significant effect via conventional power analysis (i.e., pwr.t.test), and $TP_E = 1 \times \beta$. For *"pilot"*, sample size was calculated to obtain adequate power for the mean of the $k_b^+$ positive-signed effects in the basis study (no significance-based selection), and $TP_E = (k_b^+/2) \times \beta$. The main study was then conducted, true positives defined as detections matching the ground truth direction ($x_{1|1}$, following notation in [2]), and researcher expectation errors evaluated as follows:

$$Proportion\ expected\ detects = \frac{\mathrm{TP_E}}{\mathrm{TP_{actual}}}$$

$$Spatial\ overlap = \frac{|X_{1|\cdot,m}\ \cap X_{1|\cdot,b}\ |}{|X_{1|\cdot,b}|}$$

$$Type\ M\ Error = \frac{|d_m|}{d_{true}}$$ [31]

$$Type\ S\ Error = sign(d_m)! \neq sign(d_{true})$$ [31]

For the strategy using corrected effects, $TP_E = Average\ Power(p'_{FDR}) \times k$, where average power is defined above and $k$ is the number of regions. The main study was run at the *a priori* sample size, with true positives and expectation errors defined as above. Only spatial overlap is undefined since no *a priori* spatial location is specified.

This produced 33,600 total simulated studies (200 repetitions × 4 outcome categories × 6 sample sizes × 7 study types reflecting 3 basis, 4 main).

*Effect size by proportion of the brain*

The same procedure used previously to estimate multivariate effect sizes was again used for this analysis (i.e., PCA followed by CCA or Hotelling's T-squared test), with the exception that first the largest *k* percent absolute value effect size predictors were retained before PCA was performed, across values of *k* = 1, 5, 10, 25, 50, 75, and 100 percent of the brain. Then multivariate test statistics were obtained and converted to *d* as described in previous sections. This reflects the size of the effect due to all *k* effects in combination.

### Implementation of effect size estimation procedures

*R* was used for effect size and uncertainty estimation to leverage its extensive statistical functionality, as well as to subsequently take advantage of the popular *Shiny* web app building framework used for study-level visualization (below). Built-in and custom functions as well as standard scientific libraries (e.g., *ggplot2*) were used for all above procedures with notable exceptions in the text (e.g., *metafor* for meta-analysis).

All code has been made publicly available on https://github.com/neuroprismlab . Mass univariate effect sizes and uncertainty estimation procedures are implemented in the *calculate_effeX* repository. The estimation of the cross-brain distribution of effect sizes, simulation studies, and missed effect analyses are implemented in the *crossbrain_effects* repository. Most analyses are contained within the aforementioned repositories, although the meta-analysis of spatial maps is implemented in the *BrainEffeX_utils* repository (also available as an *R* package).

### Visualization

The *BrainEffeX_utils R* package was used for visualization of all study-level results and meta-analysis of spatial maps results (Shearer et al., 2025). User- and developer-oriented details regarding usage, architecture of the accompanying *BrainEffeX R Shiny* app and utils, data structure for contributed subject-level data/meta-data, data structure for group-level effect maps/meta-data, and directions for contribution are described in (Shearer et al., 2025). In brief, the utils package facilitates ingestion and plotting of study-level effects and meta-analysis of spatial maps results for local saving or exploration through the web app.

The *effex_manuscript* repository was used for visualization of all results relating to or emerging from the estimation of the distribution of effect sizes across the brain. This repository follows the data structure conventions used in *BrainEffeX*.
