## Supplementary Tables & Figures for "The Hidden Landscape of Missed Effects in Human Functional Neuroimaging"

| Study Name (dataset_map_test_task_variables) | Dataset | N | In-Scanner Task | Outcome Measure | Measure Info |
|---|---|---|---|---|---|
| abcd_fc_r_REST_age_months | abcd | 8326 | REST | age_months | Age in months |
| abcd_fc_r_REST_bmi | abcd | 8320 | REST | bmi | Body Mass Index |
| abcd_fc_r_REST_cbcl_scr_syn_aggressive_t_FU1 | abcd | 7899 | REST | cbcl_scr_syn_aggressive_t_FU1 | Aggressive CBCL Syndrome Scale (t-score) at follow-up 1 |
| abcd_fc_r_REST_cbcl_scr_syn_aggressive_t | abcd | 8362 | REST | cbcl_scr_syn_aggressive_t | Aggressive CBCL Syndrome Scale (t-score) |
| abcd_fc_r_REST_cbcl_scr_syn_anxdep_t_FU1 | abcd | 7899 | REST | cbcl_scr_syn_anxdep_t_FU1 | AnxDep CBCL Syndrome Scale (t-score) |
| abcd_fc_r_REST_cbcl_scr_syn_attention_t_FU1 | abcd | 7899 | REST | cbcl_scr_syn_attention_t_FU1 | Attention CBCL Syndrome Scale (t-score) |
| abcd_fc_r_REST_cbcl_scr_syn_attention_t | abcd | 8362 | REST | cbcl_scr_syn_attention_t | Attention CBCL Syndrome Scale (t-score) |
| abcd_fc_r_REST_cbcl_scr_syn_external_t_FU1 | abcd | 7899 | REST | cbcl_scr_syn_external_t_FU1 | External CBCL Syndrome Scale (t-score) |
| abcd_fc_r_REST_cbcl_scr_syn_external_t | abcd | 8362 | REST | cbcl_scr_syn_external_t | External CBCL Syndrome Scale (t-score) |
| abcd_fc_r_REST_cbcl_scr_syn_internal_t_FU1 | abcd | 7899 | REST | cbcl_scr_syn_internal_t_FU1 | Internal CBCL Syndrome Scale (t-score) |
| abcd_fc_r_REST_cbcl_scr_syn_internal_t | abcd | 8362 | REST | cbcl_scr_syn_internal_t | Internal CBCL Syndrome Scale (t-score) |
| abcd_fc_r_REST_cbcl_scr_syn_rulebreak_t_FU1 | abcd | 7899 | REST | cbcl_scr_syn_rulebreak_t_FU1 | RuleBreak CBCL Syndrome Scale (t-score) |
| abcd_fc_r_REST_cbcl_scr_syn_rulebreak_t | abcd | 8362 | REST | cbcl_scr_syn_rulebreak_t | RuleBreak CBCL Syndrome Scale (t-score) |
| abcd_fc_r_REST_cbcl_scr_syn_social_t_FU1 | abcd | 7899 | REST | cbcl_scr_syn_social_t_FU1 | Social CBCL Syndrome Scale (t-score) at follow-up 1 |
| abcd_fc_r_REST_cbcl_scr_syn_somatic_t_FU1 | abcd | 7899 | REST | cbcl_scr_syn_somatic_t_FU1 | Somatic CBCL Syndrome Scale (t-score) |
| abcd_fc_r_REST_cbcl_scr_syn_thought_t_FU1 | abcd | 7899 | REST | cbcl_scr_syn_thought_t_FU1 | Thought CBCL Syndrome Scale (t-score) |
| abcd_fc_r_REST_cbcl_scr_syn_thought_t | abcd | 8362 | REST | cbcl_scr_syn_thought_t | Thought CBCL Syndrome Scale (t-score) |
| abcd_fc_r_REST_cbcl_scr_syn_withdep_t_FU1 | abcd | 7899 | REST | cbcl_scr_syn_withdep_t_FU1 | WithDep CBCL Syndrome Scale (t-score) |
| abcd_fc_r_REST_nihtbx_fluidcomp_fc | abcd | 7713 | REST | nihtbx_fluidcomp_fc | Cognition Fluid Composite Fully-Corrected T-score |
| abcd_fc_r_rest_pea_wiscv_tss | abcd | 7628 | REST | pea_wiscv_tss | WISC-V Matrix Reasoning Total Scaled Score |
| abcd_fc_t_MID_REST | abcd | 7496 | MID, REST | task - rest | |
| abcd_fc_t_nBack_REST | abcd | 7360 | nBack, REST | task - rest | |
| abcd_fc_t_SST_REST | abcd | 7417 | SST, REST | task - rest | |
| abcd_fc_t2_REST_sex | abcd | 8363 | REST | sex | |
| hbn_fc_r_rest_nih_card | hbn | 449 | REST | card | Dimensional Change Card Sort Age 3+ Age-adjusted Scale Score |
| hbn_fc_r_rest_nih_flanker | hbn | 449 | REST | flanker | Flanker Inhibitory Control and Attention Age 3+ Age-adjusted Score |

| | | | | | |
|---|---|---|---|---|---|
| hbn_fc_r_rest_nih_list | hbn | 449 | REST | list | List Sorting Working Memory Age 7+ Age-adjusted Score |
| hbn_fc_r_rest_nih_process | hbn | 449 | REST | process | Pattern Comparison Process Speed 7+ Age-adjusted Score |
| hbn_fc_r_rest_srs_srs | hbn | 641 | REST | srs | Social Responsiveness Scale |
| hbn_fc_t2_rest_diag_diag | hbn | 445 | REST | diag | Diagnosis |
| hcp_act_t_EMOTION | hcp | 314 | EMOTION | task activation | |
| hcp_act_t_GAMBLING | hcp | 324 | GAMBLING | task activation | |
| hcp_act_t_RELATIONAL | hcp | 314 | RELATIONAL | task activation | |
| hcp_act_t_SOCIAL | hcp | 317 | SOCIAL | task activation | |
| hcp_act_t_WM | hcp | 324 | WM | task activation | |
| hcp_fc_r_REST_PMAT24_A_CR | hcp | 1060 | REST | PMAT24_A_CR | Fluid Intelligence (Penn Progressive Matrices): number of correct responses |
| hcp_fc_r_REST_ProcSpeed_AgeAdj | hcp | 1067 | REST | ProcSpeed_AgeAdj | Processing Speed (Pattern Completion Processing Speed): age adjusted scale score |
| hcp_fc_t_REST_EMOTION | hcp | 1023 | EMOTION, REST | task - rest | HCP Emotion task |
| hcp_fc_t_REST_GAMBLING | hcp | 1058 | GAMBLING, REST | task - rest | HCP Gambling task |
| hcp_fc_t_REST_RELATIONAL | hcp | 1017 | RELATIONAL, REST | task - rest | HCP Relational task |
| hcp_fc_t_REST_SOCIAL | hcp | 1028 | SOCIAL, REST | task - rest | HCP Social task |
| hcp_fc_t_REST_WM | hcp | 1059 | WM, REST | task - rest | HCP Working memory task |
| hcp_fc_t2_REST_Gender | hcp | 1067 | REST, REST | Gender | |
| pnc_fc_r_REST_ADD011 | pnc | 1248 | REST | ADD011 | Attention Deficit Disorder: Did you often have trouble paying attention or keeping your mind on your school, work, chores, or other activities that you were doing? |
| pnc_fc_r_REST_DEP001 | pnc | 1251 | REST | DEP001 | Depression: Has there ever been a time when you felt sad or depressed most of the time? |
| pnc_fc_r_REST_EAT001 | pnc | 1248 | REST | EAT001 | Eating Disorder: Was there ever a time when you felt really fat or heavy, but other people said that you were too thin? |
| pnc_fc_r_REST_GAD001 | pnc | 1248 | REST | GAD001 | Generalized Anxiety Disorder: Have you ever been a worrier? |
| pnc_fc_r_REST_MAN001 | pnc | 1262 | REST | MAN001 | Mania/ Hypomania: Have there been times when you were much more active, excited or energetic than usual, had problems sitting still, or needed to move around a lot? |

| | | | | | |
|---|---|---|---|---|---|
| pnc_fc_r_REST_OCD018 | pnc | 1262 | REST | OCD018 | Obsessive Compulsive Disorder: Have you ever saved up so many things that people complained or they got in the way? |
| pnc_fc_r_REST_PAN001 | pnc | 1252 | REST | PAN001 | Panic Disorder: Have you ever had an attack like this? |
| pnc_fc_r_REST_PSY001 | pnc | 1246 | REST | PSY001 | Psychosis: Have you ever heard voices when no one was there? |
| pnc_fc_r_REST_SIP003 | pnc | 1247 | REST | SIP003 | SIPS - PRIME SCREEN-REVISED: I think that I have felt that there are odd or unusual things going on that I can't explain. |
| pnc_fc_t2_REST_Sex | pnc | 1267 | REST | Sex | Sex of participant. 0: unknown, 1: female, 2: male |
| slim_fc_r_REST_State_Anxiety | slim | 473 | REST | State_Anxiety | State anxiety measured with the State Trait Anxiety Inventory (STAI) |
| slim_fc_t2_REST_Sex | slim | 544 | REST | Sex | Participant sex |
| ukb_fc_r_rest_age | ukb | 40925 | REST | age | Age in years |
| ukb_fc_r_rest_fluid_intelligence | ukb | 37730 | REST | fluid_intelligence | Fluid intelligence [Field ID: 20016]: verbal and numerical reasoning multiple-choice questions answered correctly in two minutes (total=13). |
| ukb_fc_t2_rest_gender | ukb | 40925 | REST | gender | Participant gender |

**SI Table 1. Dataset characteristics for each study**. Dataset, sample size, neuroimaging measures, and outcome measures are provided for each study. Outcome measures encompass demographic variables, cognitive assessments, clinically relevant assessments, psychiatric screening questions, and task performance metrics across multiple cognitive domains.

| Dataset | Task | Task Info | Scan Duration |
|---|---|---|---|
| abcd | REST | Eyes open. Participants were instructed to relax while maintaining passive visual fixation on a crosshair presented centrally on screen. | 20 minutes total, with only 10 minutes used by the contributor for FC calculations |
| abcd | MID | Participants view incentive cues indicating potential monetary outcomes (Win $0.20, Win $5.00, Lose $0.20, Lose $5.00, or neutral $0), followed by a variable delay period, then a brief target stimulus (150-500ms) requiring a speeded button press response. Must always respond regardless of trial type with fast responses on win trials to earn money, fast responses on loss trials to avoid losing money, while slow responses result in no monetary gain or loss of money, respectively. Task difficulty adjusted to maintain approximately 60% success rate across all participants. | 20 minutes rest scan with only 10 minutes used by the contributor for FC calculation and 5.7 minutes task scan |
| abcd | nBack | The Emotional N-back (EN-back) task is a working memory paradigm that measures both cognitive control and emotion regulation. The task consists of two runs with eight blocks each, where participants view sequences of happy faces, fearful faces, neutral faces, and places for 2 seconds per stimulus, followed by 500ms fixation periods. In the 0-back condition (low memory load), participants respond "match" when the current stimulus matches a target image shown at the beginning of the block, essentially testing simple recognition. In the 2-back condition (high memory load), participants must respond "match" when the current stimulus is identical to the one presented two trials earlier. Each block contains 10 trials with 2 targets, 2-3 non-target lures, and several non-lures, totaling 160 trials across both runs with 80 trials per memory condition and 40 trials per stimulus type. [1,2] | 20 minutes rest scan with only 10 minutes used by the contributor for FC calculation and 5.7 minutes task scan |

| | | | |
|---|---|---|---|
| abcd | SST | Participants respond to directional arrows (left/right) as quickly as possible on "Go" trials, but must withhold their response when an unpredictable "Stop" signal (upward arrow) appears. Each of 2 runs contains 180 trials (150 Go trials, 30 Stop trials) with each trial lasting 1000ms. A tracking algorithm adjusts the Stop Signal Delay (SSD) to maintain approximately 50% successful inhibitions. SSD decreases time by 50ms after failed stops (easier) and increases by 50ms after successful stops (harder). Each run lasts 349 seconds total. | 20 minutes rest scan with only 10 minutes used by the contributor for FC calculation and 5.7 minutes task scan |
| hbn | REST | Exact quote: The participant was presented a white fixation cross in the center of a black screen and instructed to rest with eyes open. Specific instructions were as follows: "Please lie quietly with your eyes open, and direct your gaze towards the plus symbol. During this scan let your mind wander. If you notice yourself focusing on a particular stream of thoughts, let your mind wander away." (O'Connor et al. 2017) | Initially 10 minutes resting state scan eventually broken into two 5 minute scans. In both cases, all 10 minutes were used for FC calculations by the contributor |
| hcp | EMOTIO N | Subjects must decide which of two faces, with either fearful or angry expressions, presented on the bottom of the screen, matches the face at the top of the screen, or which of two shapes presented at the bottom of the screen matches the shape at the top of the screen. The contrast used in the analysis is comparing activation during face-matching blocks vs shape-matching blocks. | Each run contains 6 blocks (3 face + 3 shape) at 21 seconds per block plus 8 seconds of final fixation, totaling 134 seconds (~2.2 minutes) per run, with 2 runs making the entire emotion task approximately 4.5 minutes long. |
| hcp | GAMBLI NG | Participants play a card guessing game where they predict whether a mystery card number (1-9) is higher or lower than 5, receiving feedback in the form of monetary rewards (green arrow, +$1), losses (red arrow, -$0.50), or neutral outcomes (gray arrow, number 5). Each trial consists of a decision phase (up to 1500 ms), feedback (1000 ms), and fixation (1000 ms). The contrast is biased toward mostly rewards vs biased towards mostly losses. Each of two runs contains 2 reward blocks, 2 loss blocks, and 4 fixation blocks (15 seconds each). | Each run contains 4 task blocks (8 trials each at 3.5 seconds per trial) plus 4 fixation blocks (15 seconds each), totaling approximately 2.9 minutes per run, with 2 runs making the entire gambling task about 5.8 minutes long. |
| hcp | RELATIO NAL | Participants view shapes with different textures and perform two types of tasks: identifying whether two pairs of objects differ along shape or texture, and determining if a bottom object matches top objects on a specified shape or texture indicated by a word cue. The relational condition uses 4 trials per block (3500 ms stimulus, 500 ms ITI) while the matching condition uses 5 trials per block (2800 ms stimulus, 400 ms ITI), with both block types lasting 18 seconds total. The contrast is relational vs matching. | Each run contains 6 task blocks (18 seconds each) plus 3 fixation blocks (16 seconds each), totaling approximately 2.6 minutes, with 2 runs making the entire relational processing task about 5.2 minutes long. |
| hcp | SOCIAL | Subjects are presented with short video clips of squares, circles, and triangles that either interact in some way or move randomly on the screen. Subjects must then judge whether the objects had an interaction that appears as if the shapes are taking into account each other's feelings and thoughts; if they are not sure, or if there is no obvious interaction between the shapes, and the movement appears random. The contrast is mental interactions vs random interactions. | Each run contains 5 video blocks (20 seconds each) plus 5 fixation blocks (15 seconds each), totaling approximately 2.9 minutes per run, with 2 runs making the entire social cognition task about 5.8 minutes long. |
| hcp | WM | Subjects view images from four categories (faces, places, tools, body parts) while performing either a low-demand task, where they match the current image to a cue (0-back task), or a high-demand task where they must indicate if a current image matches the one from 2 trials ago (2-back). The contrast is faces vs the average of all other categories (places, tools, body parts) across both 0-back and 2-back conditions. | Each run contains 8 task blocks (25 seconds each) plus 4 fixation blocks (15 seconds each), totaling approximately 4.3 minutes per run, with 2 runs making the entire working memory task about 8.6 minutes long. |
| hcp | REST | Eyes open with relaxed fixation on bright crosshair against dark background in darkened room. Phase encoding alternated between right-to-left (RL) and left-to-right (LR) within sessions. | 2 sessions with 2 runs each session with 15 minutes per run. Total of 60 minutes with only 26 minutes used by the contributor for FC calculations. |
| pnc | REST | Exact quote: "During the resting-state scan, a fixation cross was displayed as images were acquired. Subjects were instructed to stay awake, keep their eyes open, fixate on the displayed crosshair, and remain still." (Satterthwaite et al. 2013) | 6.18 minutes |
| slim | REST | Exact quote: "During the resting-state MRI scan, the subjects were instructed to lie down, close their eyes, and rest without thinking about any specific thing but to refrain from falling asleep." (Wei et al. 2018) | 8 minutes |

| | | | |
|---|---|---|---|
| ukb | REST | Exact quote: "Subjects are instructed to keep their eyes fixated on a crosshair, relax and “think of nothing in particular” (Miller et al. 2018) | 6.17 minutes |

**SI Table 2. Description of fMRI tasks and scan durations across all datasets.** The experimental protocol, stimulus presentation details, and scan duration are provided for each dataset and task. Scan durations indicate total acquisition time and, where applicable, the portion used for functional connectivity (FC) calculations. Resting-state scans involved passive fixation with eyes open (ABCD, HBN, HCP, PNC, UKB) or closed (SLIM), ranging from 6-60 minutes. Task-based paradigms describe trial structure, timing parameters, and relevant contrasts used in analyses. Tasks included monetary incentive delay (MID), emotional n-back working memory (nBack), stop-signal inhibition (SST), emotion face matching (EMOTION), card gambling (GAMBLING), relational reasoning (RELATIONAL), social cognition with geometric animations (SOCIAL), and category-based working memory (WM). Task durations ranged from 2.2-8.6 minutes. [1, 2, 3, 4, 5, 6, 7, 8, 9, 10].

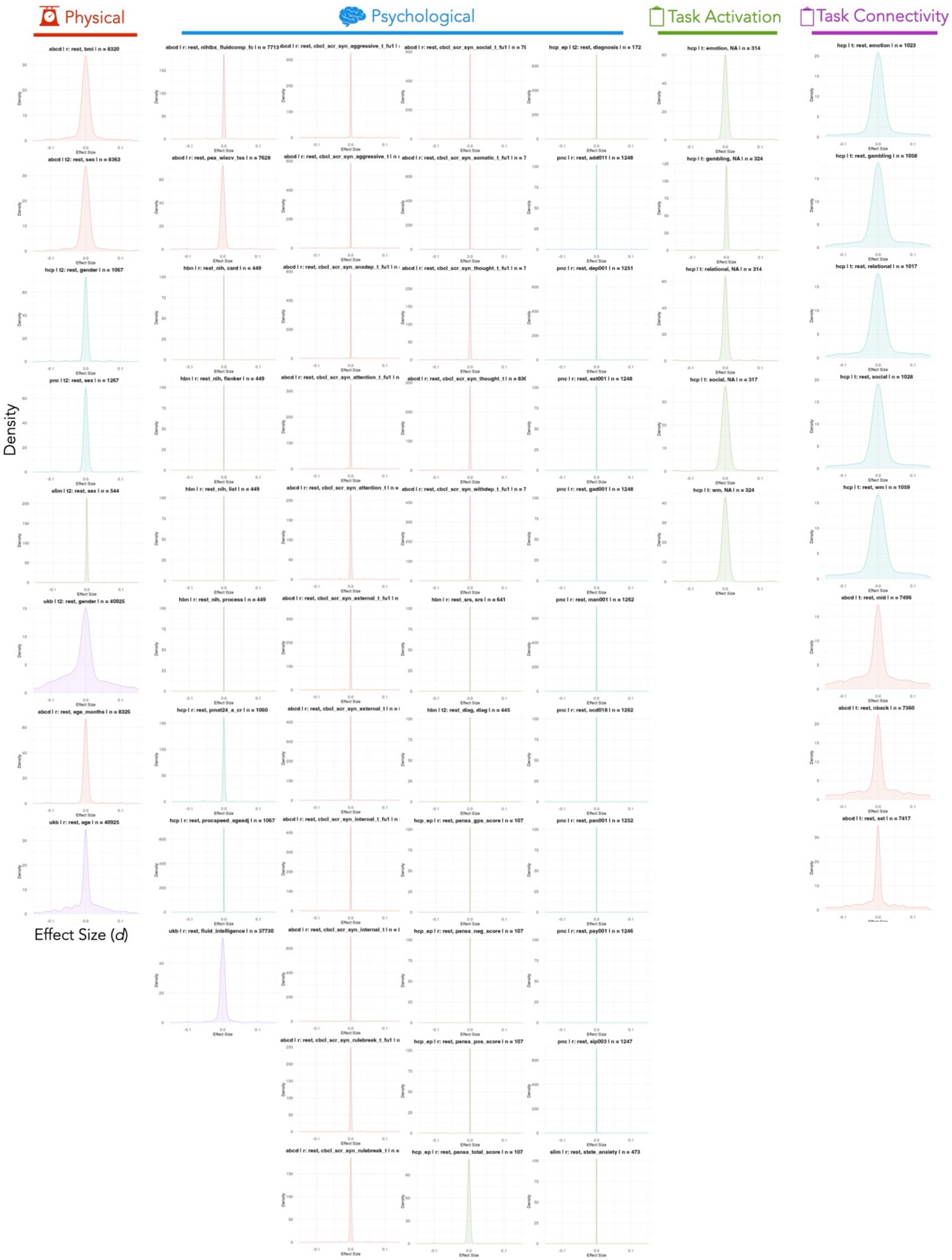


**SI Figure 1. Uncorrected mass univariate cross-brain effect size distributions for all studies.** Density plots show the distribution of Cohen's d values across edges or voxels in each study.

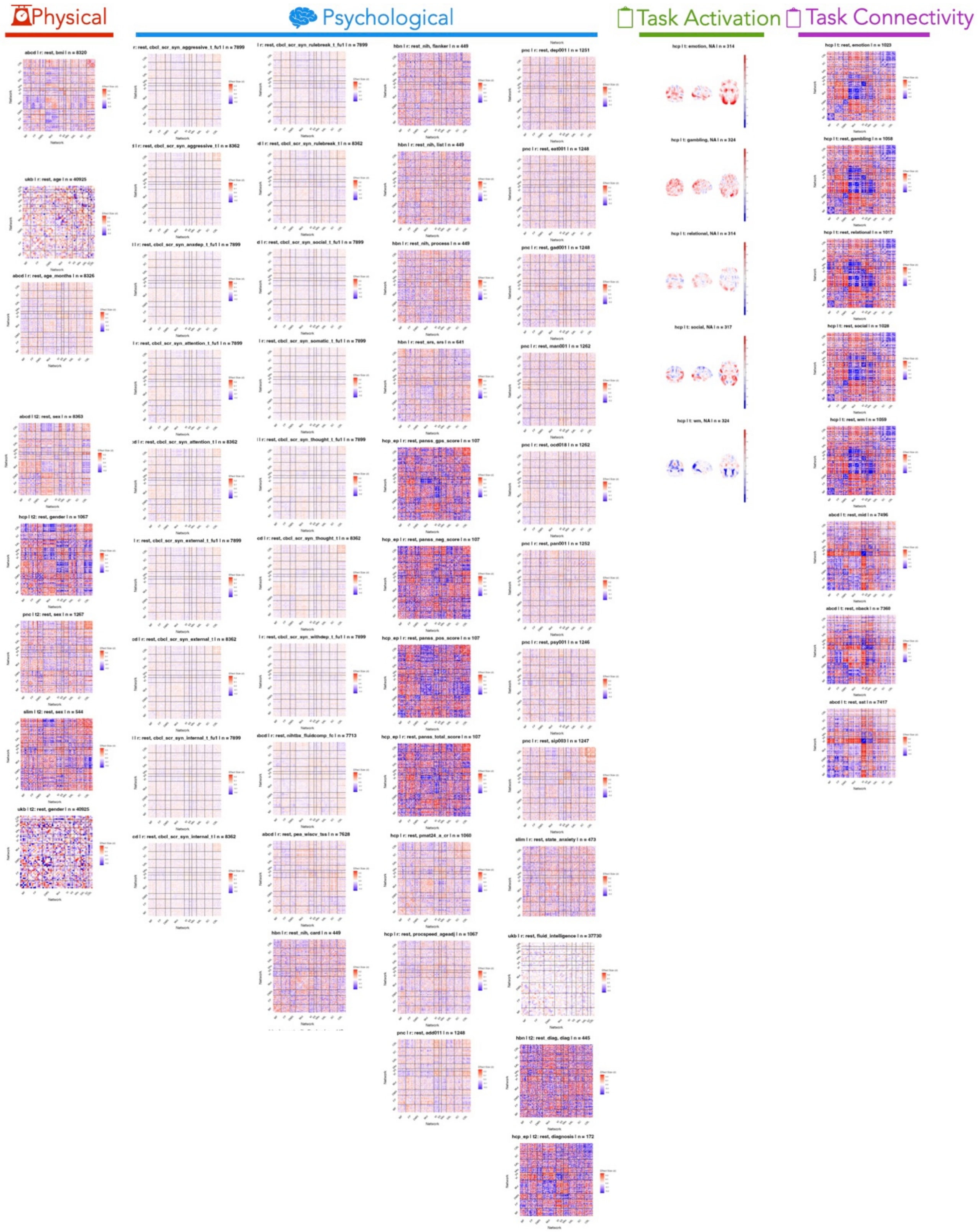


**SI Figure 2. Spatial maps of effect size point estimates for all studies**. Functional connectivity maps are arranged by network and activation maps are arranged spatially.

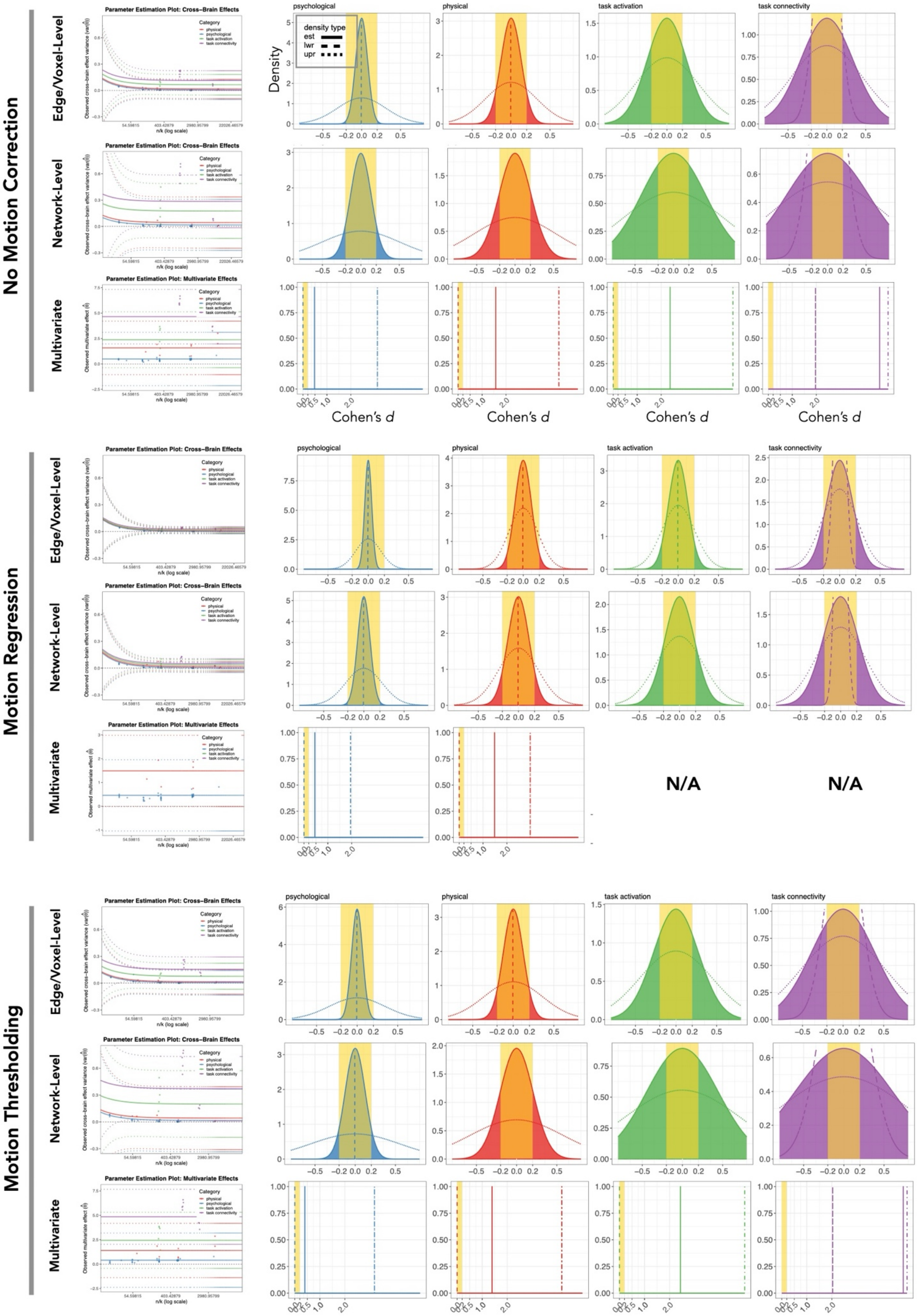


**SI Figure 3. Cross-brain effect size distribution by motion deconfounding strategy.** Parameter estimation plots and resulting corrected cross-brain effect size distributions for no control, motion regression, and motion thresholding.

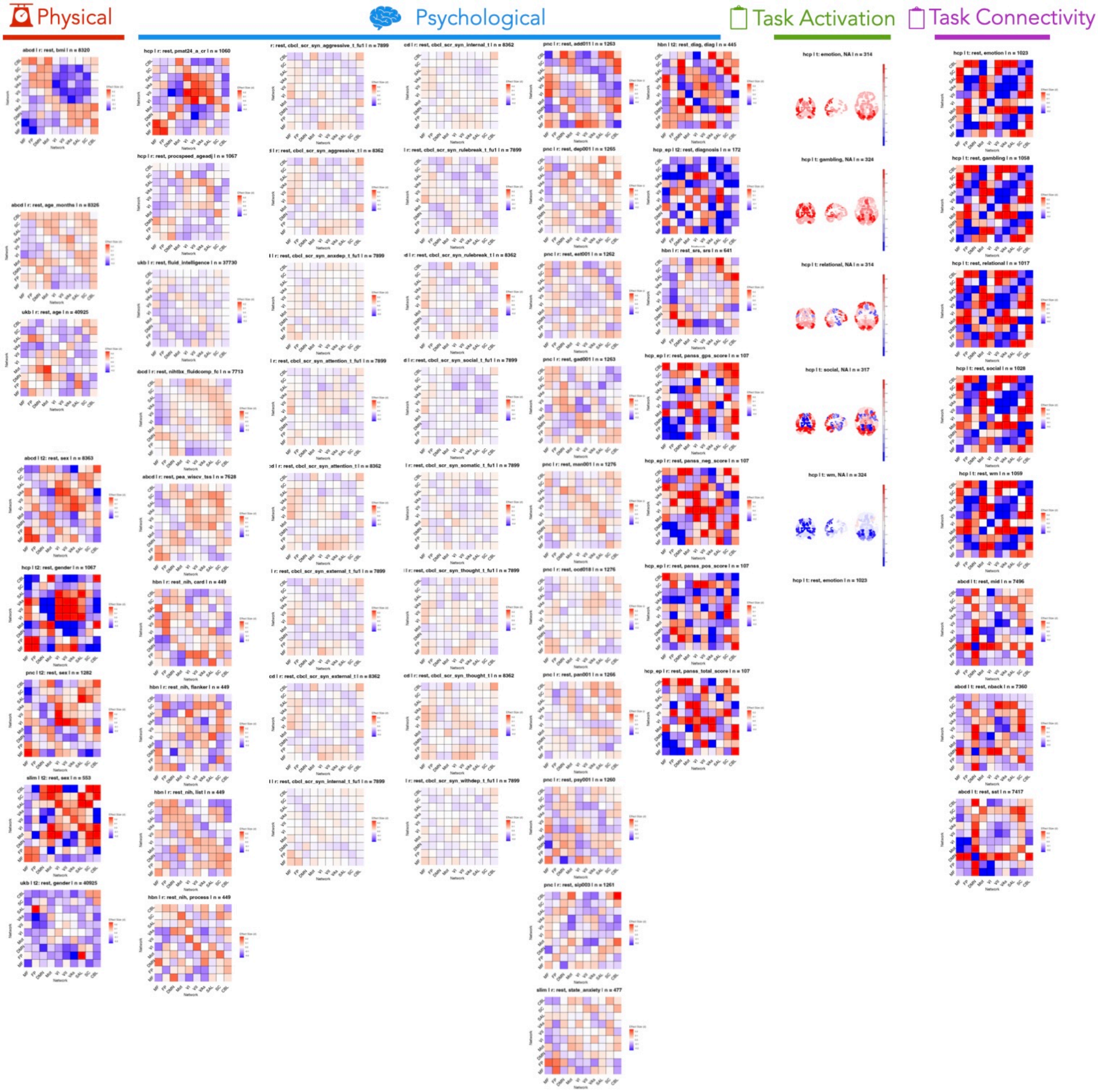


**SI Figure 4. Spatial maps of network-level effect size point estimates for all studies.** Functional connectivity is arranged by network, and activation maps are arranged spatially.

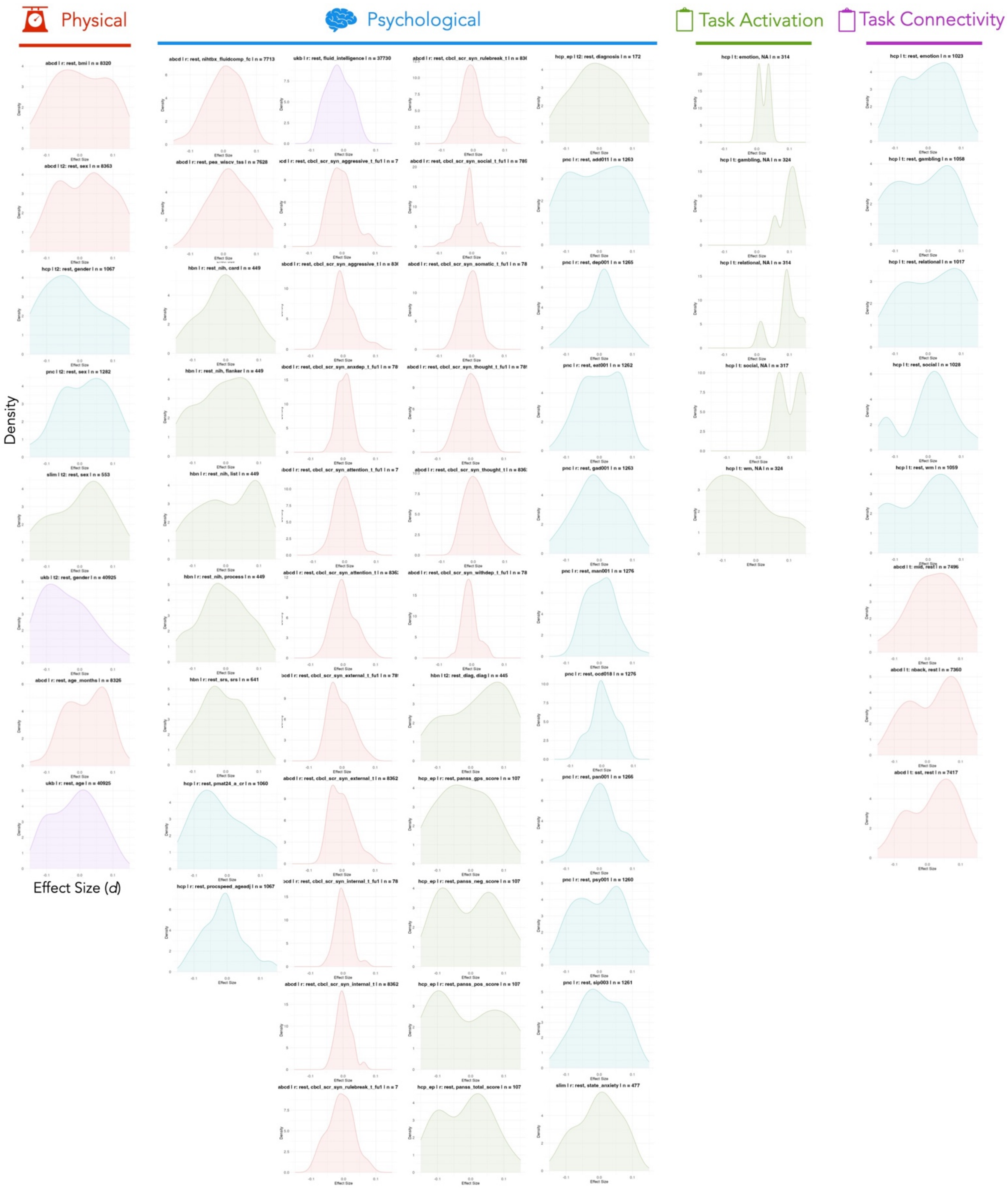


**SI Figure 5. Network-level uncorrected mass univariate cross-brain effect size distributions for all studies.** Density plots show the distribution of Cohen's d values across networks in each study.

| Edge-/voxel-level tau | | | |
|---|---|---|---|
| | *lwr* | *est* | *upr* |
| *psychological* | 0 | 0.04319044 | 0.15521454 |
| *physical* | 0 | 0.10182351 | 0.1816839 |
| *task activation* | 0 | 0.12011973 | 0.20485957 |
| *task connectivity* | 0.06399325 | 0.16350695 | 0.22220241 |

| Network-level tau | | | |
|---|---|---|---|
| | *lwr* | *est* | *upr* |
| *psychological* | 0 | 0.07698691 | 0.22552539 |
| *physical* | 0 | 0.13238866 | 0.25126588 |
| *task activation* | 0 | 0.1851588 | 0.2912742 |
| *task connectivity* | 0.05719886 | 0.2222323 | 0.30903508 |

| Multivariate point estimates | | | |
|---|---|---|---|
| | *lwr* | *est* | *upr* |
| *psychological* | -1.0406585 | 0.45715696 | 1.95497247 |
| *physical* | -0.0129068 | 1.48490873 | 2.98272423 |
| *task connectivity* | NA | NA | NA |
| *task activation* | NA | NA | NA |

| Multivariate point estimates (0.1mm FFD thresholding) | | | |
|---|---|---|---|
| | *lwr* | *est* | *upr* |
| *psychological* | -2.3735835 | 0.40348583 | 3.18055514 |
| *physical* | -1.3894623 | 1.39451864 | 4.17849958 |
| *task connectivity* | 2.02858739 | 4.84731202 | 7.66603665 |
| *task activation* | -0.4381661 | 2.43596962 | 5.31010529 |

**SI Table 3.** Parameter estimates for effect size distributions: tau for edge-/voxel-level and network-level effect sizes, and point estimates for multivariate effect sizes.

| Edge-/voxel-level proportion | | | |
|---|---|---|---|
| | $|d|<0.1$ | $|d|<0.2$ | $|d|<0.5$ |
| *psychological* | 0.987 | >0.999 | 1.000 |
| *physical* | 0.683 | 0.954 | >0.999 |
| *task activation* | 0.595 | 0.904 | >0.999 |
| *task connectivity* | 0.468 | 0.789 | 0.998 |

| Network-level proportion | | | |
|---|---|---|---|
| | $|d|<0.1$ | $|d|<0.2$ | $|d|<0.5$ |
| *psychological* | 0.789 | 0.988 | 1.000 |
| *physical* | 0.558 | 0.876 | >0.999 |
| *task activation* | 0.401 | 0.707 | 0.992 |
| *task connectivity* | 0.351 | 0.637 | 0.977 |

| | Multivariate results | | |
|---|---|---|---|
| | *\|d\|<0.1* | *\|d\|<0.2* | *\|d\|<0.5* |
| *psychological* | 0 | 0 | 1 |
| *physical* | 0 | 0 | 0 |
| *task connectivity* | NA | NA | NA |
| *task activation* | NA | NA | NA |

| | Multivariate results (0.1mm FFD thresholding) | | |
|---|---|---|---|
| | *\|d\|<0.1* | *\|d\|<0.2* | *\|d\|<0.5* |
| *psychological* | 0 | 0 | 1 |
| *physical* | 0 | 0 | 0 |
| *task connectivity* | 0 | 0 | 0 |
| *task activation* | 0 | 0 | 0 |

**SI Table 4.** Proportion of effect sizes below conventional small and medium thresholds. For multivariate effects, this reflects whether the effect size point estimate is below the threshold (0=above, 1=below).

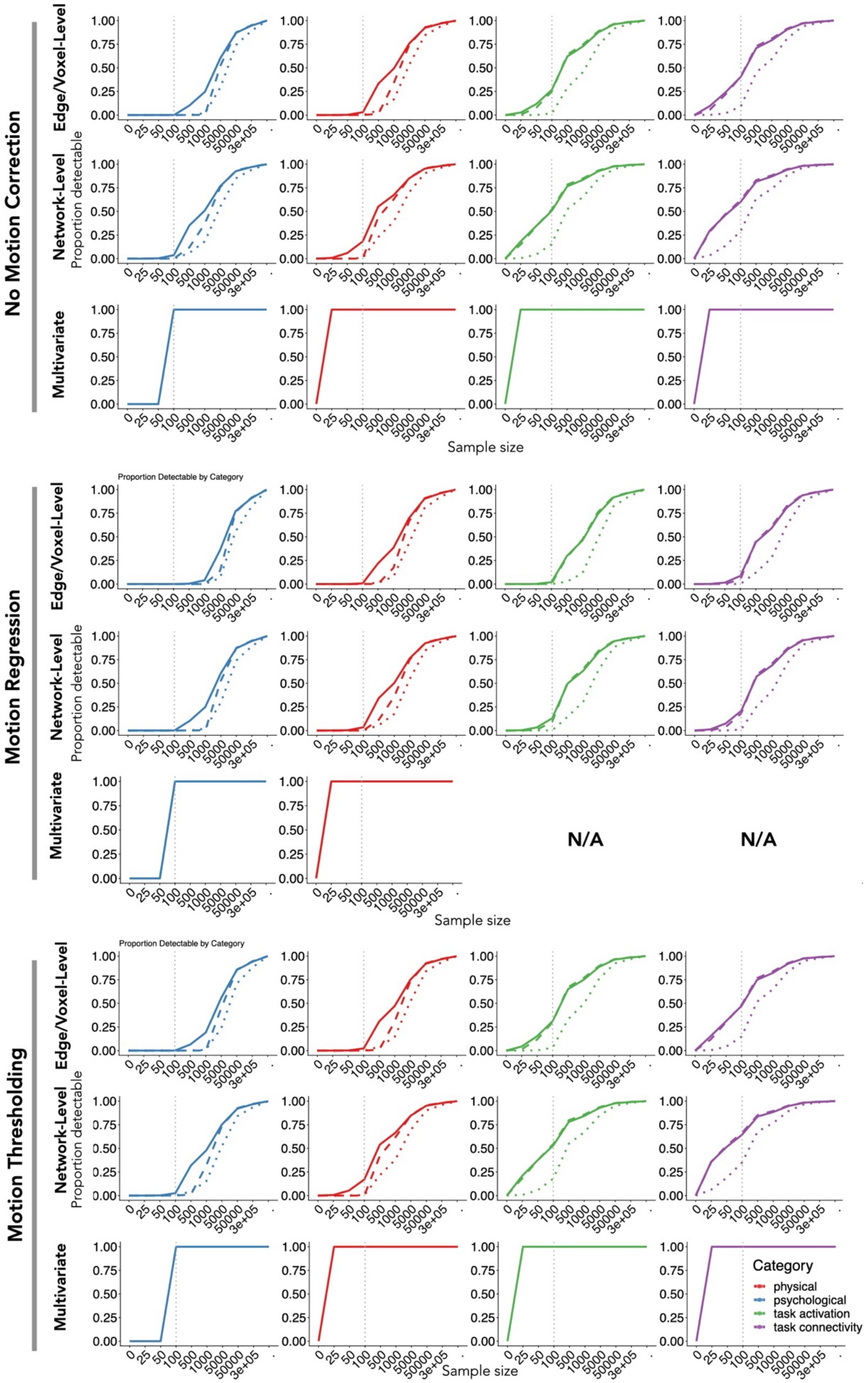


**SI Figure 6. Power analysis results by motion deconfounding strategy.** Sample size required to detect effects with 80% power for edge- or voxel-level (top), network-level (middle), or multivariate (bottom) effects for each outcome category (red and top, physical; blue and middle, psychological; green and bottom, task-based). Each plot shows the proportion of effects across the brain that can be detected with 80% power in each sample size category after Bonferroni, false discovery rate (FDR), or no correction.

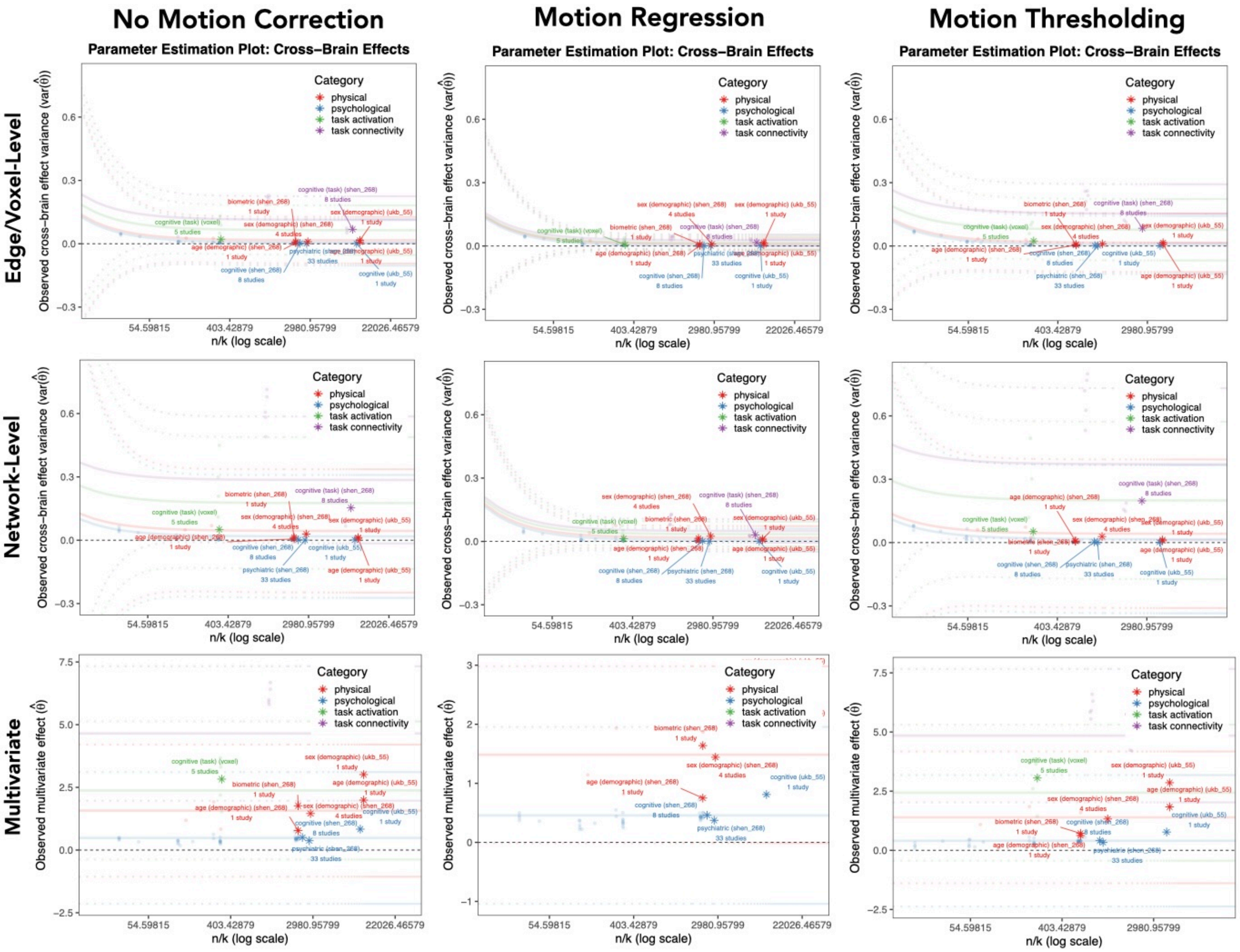


**SI Figure 7. Meta-analyses from sub-categories by motion deconfounding procedure.** $\tau_n^2$ estimation plot (transparent) overlaid with $var(\hat{\theta})$ from meta-analysis results for each sub-category (stars).

| Cohen's d | Sample size required for 80% power in within-subject design | Sample size *per group* required for 80% power in between-subject design |
|---|---|---|
| 0.2 | 198 | 393 |
| 0.5 | 33 | 64 |
| 0.8 | 14 | 26 |

**SI Table 5.** Required sample size to detect univariate within vs. between-group effects at the commonly targeted 80% power level with a common alpha threshold of 0.05.

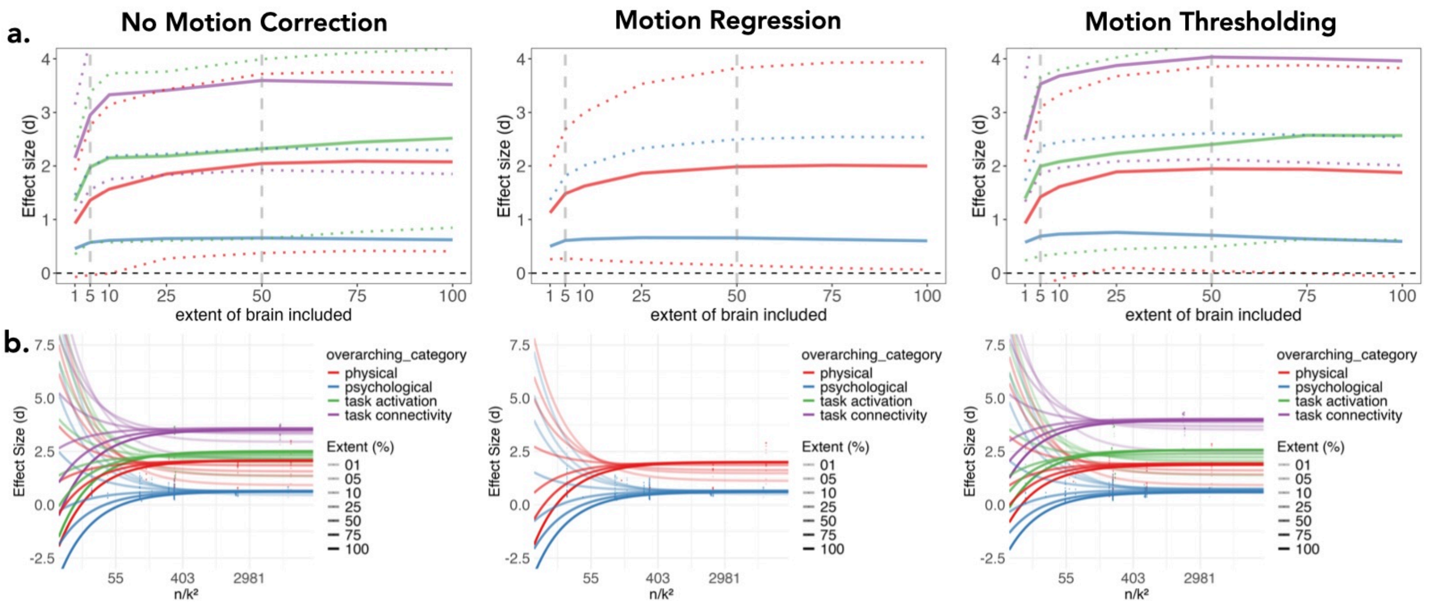


**SI Figure 8. Effect size increases with the proportion of the brain included. Top panel: a)** Multivariate effect size from the top *k* strongest effects, by outcome category (red, physical; blue, psychological; green, task activation; purple, task connectivity). The top 1%, 5%, 10%, 25%, 50%, 75%, and 100% marginal effects (x-axis) were selected and used to estimate the multivariate effect size. **b)**. The parameter estimation plot for each top *k* effect, with higher k in more opaque colors. Dotted lines denote top 5% and top 50% of effects.
